# Cortical integration of higher-order thalamic inputs is lineage-dependent

**DOI:** 10.1101/2022.03.28.486015

**Authors:** Matthew J. Buchan, Kashif Mahfooz, Joram J. van Rheede, Gemma Gothard, Sophie V. Avery, Tommas J. Ellender, Sarah E. Newey, Colin J. Akerman

## Abstract

Primary sensory cortex receives and integrates inputs from first-order and higher-order thalamic nuclei. First-order inputs convey sensory information from the periphery and exhibit simple response properties, whereas higher-order inputs exhibit more complex response properties, provide contextual feedback, and can modulate first-order inputs. Here we show that the way in which cortical neurons integrate these thalamic inputs, reflects the progenitor cell from which the cortical neurons derive. Within layer 4 of mouse primary somatosensory cortex, excitatory neurons that derive from apical intermediate progenitors exhibit multi-whisker response properties and receive higher-order thalamic input, in a manner consistent with their dendritic morphology. These properties depend upon the expression levels of the transcription factor Lhx2, which when increased, abolishes the higher-order properties of apical intermediate progenitor-derived neurons, and disrupts the induction of sensory-evoked plasticity. These data reveal a lineage-dependent mechanism that establishes the integration and functional contribution of higher-order thalamic inputs within cortex.

## Introduction

The response properties of cortical neurons reflect the thalamic inputs that they receive ^1–4^. First-order thalamic nuclei are driven by subcortical input (e.g. the retina) and are considered the primary relay from the sensory periphery to cortex, transferring sensory information via neurons with simple response properties ^5^. In contrast, higher-order thalamic nuclei receive input from multiple sources that can be of a cortical or subcortical origin ^6–9^. Consequently, neurons of higher-order nuclei display more complex response properties, reflecting the encoding of both sensory and contextual information ^6,10^. In addition to facilitating the transfer of sensory information through an ascending hierarchy of transthalamic pathways ^11,12^, higher-order thalamic nuclei also send feedback projections to lower cortical areas, where they are thought to influence the processing and plasticity of first-order input ^13–16^.

Even within layer 4 (L4), one of the earliest stages of cortical processing, there is evidence for the integration of first-order and higher-order thalamic inputs. In rodent primary somatosensory cortex (S1), tactile information from single whiskers is conveyed via neurons in the first-order, ventral posterior medial nucleus (VPM) of the thalamus, which project to discrete anatomical structures within L4, called barrels, forming a somatotopic map of the mystacial pad ^17,18^. Meanwhile, information from multiple whiskers is conveyed via neurons in the higher-order, posterior medial nucleus (POm), which receives inputs of both a cortical and subcortical origin ^6,8^. POm projections to S1 target the inter-barrel regions within L4, called septa, as well as L1 and L5a ^19–23^. Previous work, using both electrical and optical stimulation methods, has shown that whilst VPM represents the major input to L4, individual neurons differ in the degree to which they receive higher-order input from POm ^24–26^. The L4 population is therefore heterogeneous in terms of its integration of thalamic input, with some neurons receiving more higher-order information than others. However, the mechanisms that determine how individual neurons sample thalamic inputs are unknown.

One potential explanation is that the thalamic inputs to a cortical neuron are reflective of the neuron’s developmental history, or lineage. All excitatory neurons are born from a heterogeneous pool of progenitor cells, which reside in the ventricular proliferative zones during embryonic development ^27^. Previous work has shown that the local synaptic connectivity between excitatory cortical neurons reflects the progenitor from which they derive ^28–30^. Here, we examine how lineage relates to the thalamic inputs received by L4 excitatory neurons in mouse S1, by performing longitudinal studies that enable us to relate the properties of mature cortical neurons to the progenitors from which they were derived during embryonic development. Using a combination of *in utero* labelling, functional anatomy, and *in vitro* and *in vivo* electrophysiology, we characterise L4 neurons derived from a progenitor population called apical intermediate progenitors (aIPs) and compare them to L4 neurons derived from other progenitors (OPs). We find that aIP-derived neurons exhibit multi-whisker receptive fields, and preferentially receive input from higher-order thalamus compared to OP-derived neurons. Moreover, these properties are associated with low expression levels of the transcription factor Lhx2 in aIP-derived neurons, which when increased, causes aIP-derived neurons to exhibit single-whisker receptive fields, receive fewer higher-order inputs, and disrupts the induction of sensory-evoked plasticity. Thus, we describe a lineage-dependent mechanism by which different sources of thalamic information are integrated and processed within cortex.

## Results

### L4 neurons exhibit heterogeneity in their response properties

To investigate the response properties of L4 excitatory neurons in S1, we performed extracellular recordings in anaesthetised mice at postnatal day 60 (P60). Responses were measured to deflections of the principal whisker (PW) and an adjacent whisker (AW; **Figure 1A**; see **Methods**) and regular spiking (primarily excitatory) neurons were identified based on their waveform properties (Extended Data **Figure 1**). L4 was identified using the current-source density (CSD) profile following PW deflection (**Figure 1B**) and the electrode location was confirmed by *post-hoc* histological analysis (**Figure 1C**). Correct identification of the PW and AW was confirmed offline (**Figure 1D**; see **Methods**). Consistent with previous work, individual L4 excitatory neurons were responsive to PW and AW deflection ^32,33^. To quantify the response properties of L4 neurons we defined a selectivity index (SI), where higher values indicated neurons that responded mainly to the PW, and SI values closer to 0.5 indicated similar levels of response to the PW and AW - consistent with multi-whisker response properties. We assessed the SI following either single whisker deflections or trains of whisker deflections (four deflections at 10 Hz). Some L4 excitatory neurons displayed very high SI values, but others displayed SI values close to 0.5 (**Figure 1E**). Consequently, whilst L4 excitatory neurons were more responsive to deflections of the PW (**Figure 1F-G**; single deflection mean SI was 0.61 ± 0.04; train deflection mean SI was 0.64 ± 0.04), the range of SI values revealed that individual L4 excitatory neurons exhibit heterogeneity in their response properties (single deflection IQR was 0.48 to 0.76; train deflection IQR was 0.53 – 0.79).

**Figure 1:**
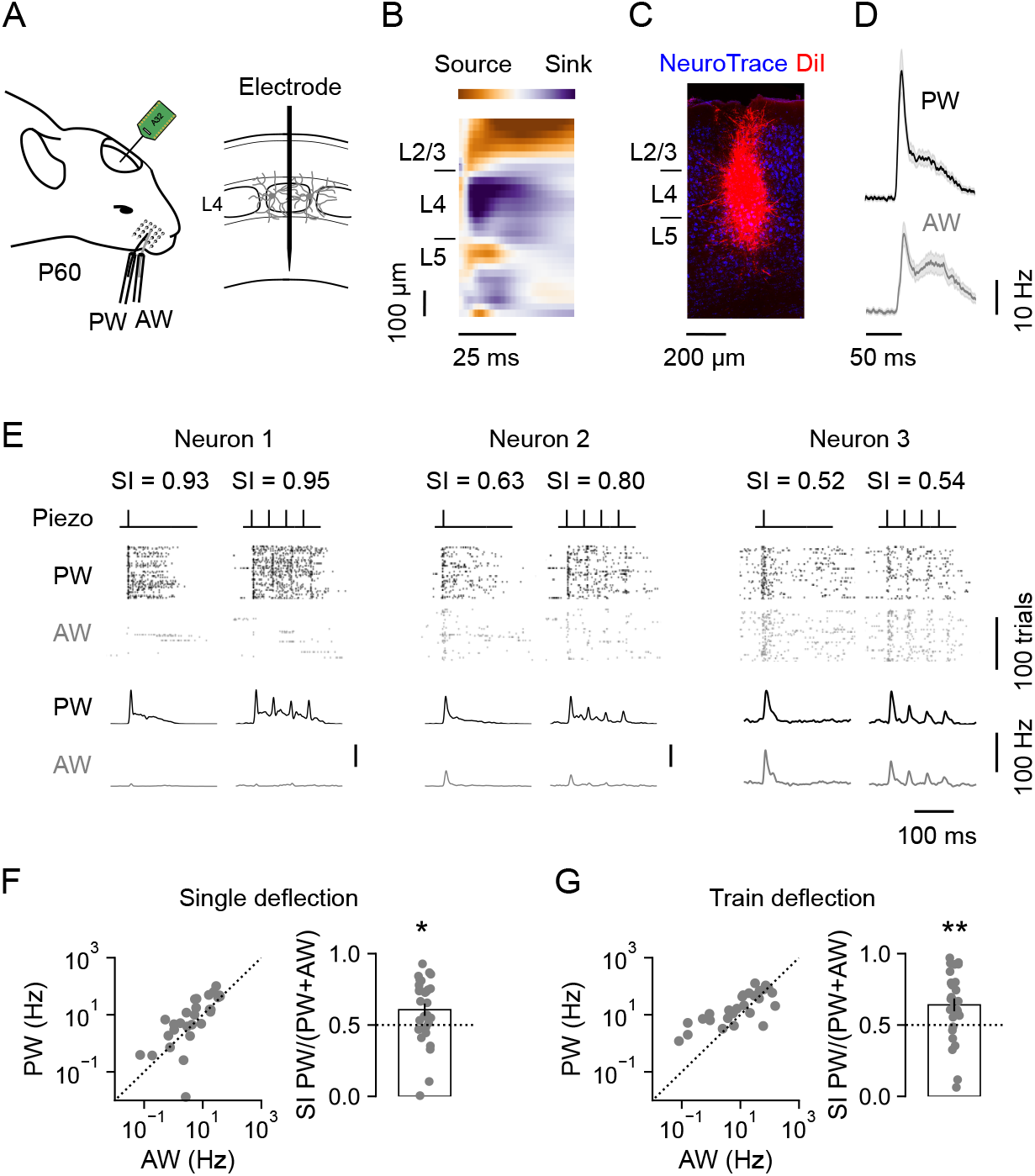
L4 neurons exhibit heterogeneous response properties in mouse S1. (A) The activity of individual regular-spiking L4 neurons in S1 was recorded in response to the deflection of the PW or AW at P60. (**B**) L4 was identified using the CSD profile following PW deflection. (**C**) Histological analysis confirmed electrode location. (**D**) Mean L4 neuronal responses from an individual animal following a single deflection of the PW (upper) or AW (lower). Shading indicates SEM around the mean. (**E**) Three L4 neurons with different SI, as defined by relative response to the PW and AW. Raster plots and corresponding PSTHs show spiking activity over 100 trials of either a single deflection (inner left) or train deflection (inner right) of the PW (black) or AW (grey). (**F**) Responses of individual neurons to single deflection of the PW and AW (left), and the distribution of corresponding SI values (right). L4 neurons were selective for the PW (n = 28 neurons; p = 0.019, one sample t-test). (**G**) Responses of individual neurons to train deflection of the PW and AW (left), and the corresponding SI values (right). L4 neurons were selective for the PW (n = 28 neurons; p = 0.004, one sample t-test). Data represented as mean ± SEM.

### Response properties of L4 neurons are lineage-dependent

To investigate whether heterogeneity in the response properties of L4 excitatory neurons is associated with lineage, we used *in utero* electroporation (IUE; see **Methods**) to pulse-label a population of dividing excitatory progenitor cells at embryonic gestation day 14 (E14), when L4 excitatory neurons are being born ^34,35^. Consistent with our and others’ previous work, we made use of the fact that the tubulin alpha1 (T*α*1) promoter can be used to label apical intermediate progenitors (aIPs) within the ventricular zone ^29,36,37^. Based upon their morphology and cell cycle behaviours, aIPs have been distinguished from other progenitors (OPs), including radial glial cells within the VZ, and outer radial glia and basal intermediate progenitor cells within the subventricular zone ^27,36,37^.

To characterise the response properties of aIP-derived and OP-derived L4 neurons in postnatal cortex, we used optotagging as an established method for identifying specific cell types *in vivo* (**Figure 2A-C**) ^38^. This involved electroporating two DNA constructs: a T*α*1-Cre construct in which Cre recombinase is under the control of a portion of the T*α*1 promoter ^37^, and a floxed ChR2-YFP reporter construct that expresses ChR2 fused to enhanced yellow fluorescent protein (YFP) following Cre recombination (**Figure 2A**). Thus, in line with our previous work, ChR2-YFP was expressed in aIP-derived L4 neurons, but not in L4 neurons derived from OPs^29^. IUE was targeted to the region of the VZ that gives rise to S1 excitatory neurons, such that when the animals reached postnatal ages, sparse labelling of aIP-derived ChR2-YFP^+^ neurons could be observed in L4 of S1 (**Figure 2B**). Neurons that exhibited reliable, short-latency (<5 ms) light-evoked responses were considered to be directly activated by ChR2 and therefore classified as aIP-derived (**Extended Data Figure 2**). Those that exhibited a longer latency, or no, light-evoked response, were classified as OP-derived.

**Figure 2:**
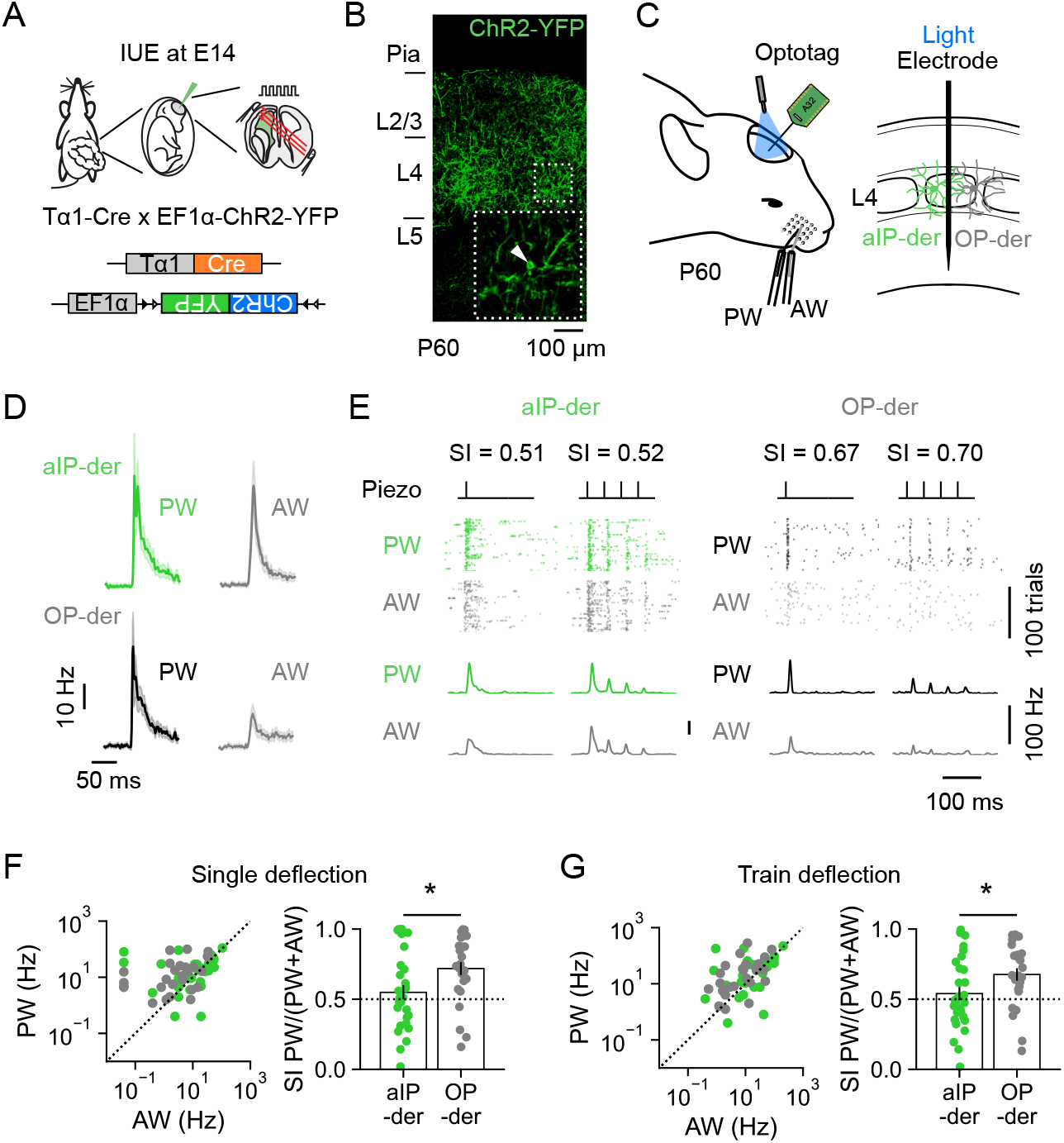
aIP-derived L4 neurons exhibit multi-whisker response properties. (**A**) IUE of a T*α*1-Cre and floxed ChR2-YFP plasmid was used to optotag aIP-derived L4 neurons in S1. (**B**) ChR2-YFP^+^ aIP-derived L4 neurons at P60. (**C**) The spiking activity of optotagged aIP-derived L4 neurons and neighbouring (non-optotagged) OP-derived L4 neurons was recorded in response to the deflection of the PW or AW. (**D**) Mean aIP-derived (top) and OP-derived (bottom) L4 responses from an individual animal following a single deflection of the PW (left) or AW (right). (**E**) Spiking of an individual aIP-derived (left, green) and OP-derived L4 neuron (right, black) over 100 trials of either single deflection (inner left) or train deflection (inner right) of the PW or AW. (**F**) Responses of individual aIP-derived and OP-derived L4 neurons to single deflection of the PW and AW (left), and the distribution of corresponding SI values (right). aIP-derived L4 neurons were less selective to the PW, and therefore relatively more responsive to the AW, when compared to OP-derived neurons (n = 29 and 26; p = 0.019, Mann Whitney U test). (**G**) Responses to train deflection of the PW and AW (left), and corresponding SI values (right). aIP-derived L4 neurons were less selective to the PW, and therefore relatively more responsive to the AW (n = 29 and 26; p = 0.040, t-test). Data represented as mean ± SEM, n = neurons, conventions as in Figure 1.

As above, deflections of the PW and AW evoked responses in both the aIP-derived and OP-derived L4 neurons (**Figure 2D**). However, compared to OP-derived neurons, aIP-derived neurons showed stronger relative responses to the AW. This was reflected in lower SI values for the aIP-derived neurons to both single and train deflections of the whiskers (**Figure 2E-G**; single deflection mean SI was 0.55 ± 0.05 for aIP-derived and 0.71 ± 0.05 for OP-derived; train deflection mean SI was 0.54 ± 0.05 for aIP-derived and 0.67 ± 0.04 for OP-derived). These data reveal that aIP-derived neurons exhibit greater multi-whisker responsivity than OP-derived neurons, suggesting that the response properties of L4 excitatory neurons reflect the progenitor type from which they derive.

### The dendritic morphology of L4 neurons is lineage-dependent

Response properties in L4 of S1 reflect a neuron’s morphology and how this relates to the organisation of thalamic inputs to barrels and septa (**Figure 3A**) ^20,21,23^. For example, L4 neurons whose soma reside outside of a barrel are more likely to exhibit multi-whisker responses ^32^. Equally, L4 neurons often target their dendrites towards the core of a single barrel, consistent with selectivity for a single, principal whisker ^39–42^. To investigate the soma position and dendritic morphology of aIP-derived and OP-derived L4 neurons, we used a second IUE labelling strategy (**Figure 3B**).

**Figure 3:**
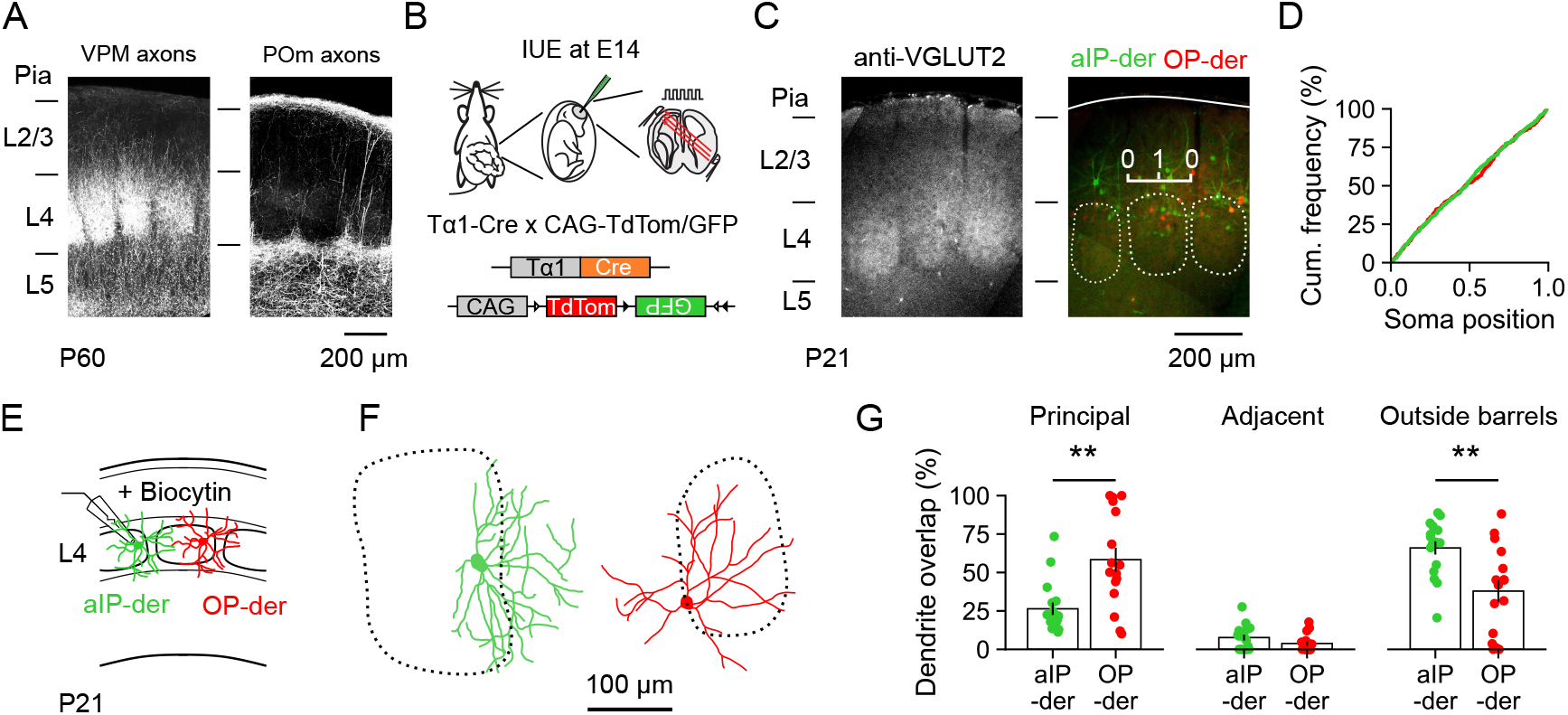
aIP-derived and OP-derived L4 neurons differ in their dendritic morphology. (**A**) Thalamic axonal input to S1 from VPM (left) and POm (right), visualised with ChR2-GFP. (**B**) IUE of a T*α*1-Cre and two-colour CAG-FLEx reporter plasmid was used to label aIP-derived (GFP^+^, green) and OP-derived (tdTomato^+^, red) L4 neurons in S1. (**C**) VGLUT2 immunohistochemistry at P21 (left) was used to relate the distribution of labelled soma to the organisation of barrels and septa. A soma position index was defined, with a value of 1 indicating a soma in the centre of a barrel (right). (**D**) There was no significant difference in the distribution of aIP-derived or OP-derived L4 neurons (n = 537 and 351; p = 0.826, t-test). (**E**) Biocytin fills were used to reconstruct the dendritic morphology of aIP-derived and OP-derived neurons. (**F**) Example aIP-derived (left) and OP-derived (right) L4 neuron reconstructions. (**G**) Dendrites of OP-derived neurons were more likely to target the principal barrel than aIP-derived neurons (left; n = 17 and 16; p = 0.005, Mann-Whitney U test). Dendrites of both populations exhibited a similar degree of overlap with the adjacent barrel (middle; n = 17 and 16; p = 0.145, Mann-Whitney U test). Dendrites of aIP-derived neurons were more likely to target the area outside of barrels, including septa (right; n = 17 and 16; p = 0.004, Mann-Whitney U test). Data represented as mean ± SEM, n = neurons.

Here, T*α*1-Cre was electroporated with a CAG-FLEx reporter construct that incorporates a flexible excision cassette, where Cre recombination permanently switches expression from tdTomato fluorescent protein to enhanced green fluorescent protein (GFP) ^43^. As such, GFP^+^ and tdTomato^+^ L4 neurons could be observed at P21, which corresponded to aIP-derived and OP-derived neurons, respectively ^29^. We defined a soma position index for the electroporated neurons, where a value of 1 indicates a soma located at the centre of a barrel, and 0 indicates the midpoint between two barrel boundaries (**Figure 3C**; see **Methods**). This index revealed that there was no significant difference in the distribution of soma position between aIP-derived and OP-derived L4 neurons at P21 (**Figure 3D**; mean soma position index was 0.48 ± 0.01 for aIP-derived and 0.48 ± 0.02 for OP-derived).

To examine dendritic morphology, we performed targeted *in vitro* whole-cell patch clamp recordings and filled individual neurons with biocytin (**Figure 3E**; see **Methods**). As reported previously, there were no differences in intrinsic electrical properties between aIP-derived and OP-derived L4 neurons, or in the frequency and amplitude of their spontaneous excitatory synaptic inputs (**Extended Data Figure 3**) ^29^. Biocytin-filled cells were reconstructed, co-registered to immunofluorescence images of the barrel field, and the overlap of the dendrite with the principal barrel was quantified (see **Methods**). The dendrites of OP-derived L4 neurons tended to target the principal barrel (**Figure 3F-G**; OP-derived overlap with principal barrel was 58.32 ± 7.78 %, adjacent barrel was 3.80 ± 1.44 %, and area outside barrels was 37.88 ± 7.19 %). In contrast, the dendrites of aIP-derived L4 neurons tended to overlap with areas outside of the principal barrel, including septa (**Figure 3F-G**, aIP-derived overlap with principal barrel was 26.38 ± 4.03 %, adjacent barrel was 7.67 ± 1.91 %, and area outside barrels was 65.96 ± 4.31 %). Although the distribution of neuronal soma is comparable for the progenitor populations characterised here, a L4 neuron’s dendritic morphology appears to reflect the neuron’s lineage. One prediction is that these differences in the morphology of the L4 neurons will influence their integration of inputs from different sources, including the thalamus ^20,21,23,32^.

### The sampling of thalamic inputs by L4 neurons is lineage-dependent

To test this prediction directly, we used the same IUE strategy at E14 to label aIP-derived and OP-derived neurons with GFP and tdTomato, respectively (**Figure 4A**). Once the animal had reached P21, a thalamic injection of an AAV expressing ChR2-GFP was stereotaxically targeted to either VPM or POm (**Figure 4A-C**; see **Methods**). This experimental design enabled us to prepare acute brain slices at P60, and record from neuronal pairs comprising a GFP^+^ aIP-derived and a tdTomato^+^ OP-derived L4 neuron, whilst using light pulses (1 ms, 473 nm) to selectively activate ChR2-expressing thalamic axons in S1 (see **Methods**).

**Figure 4:**
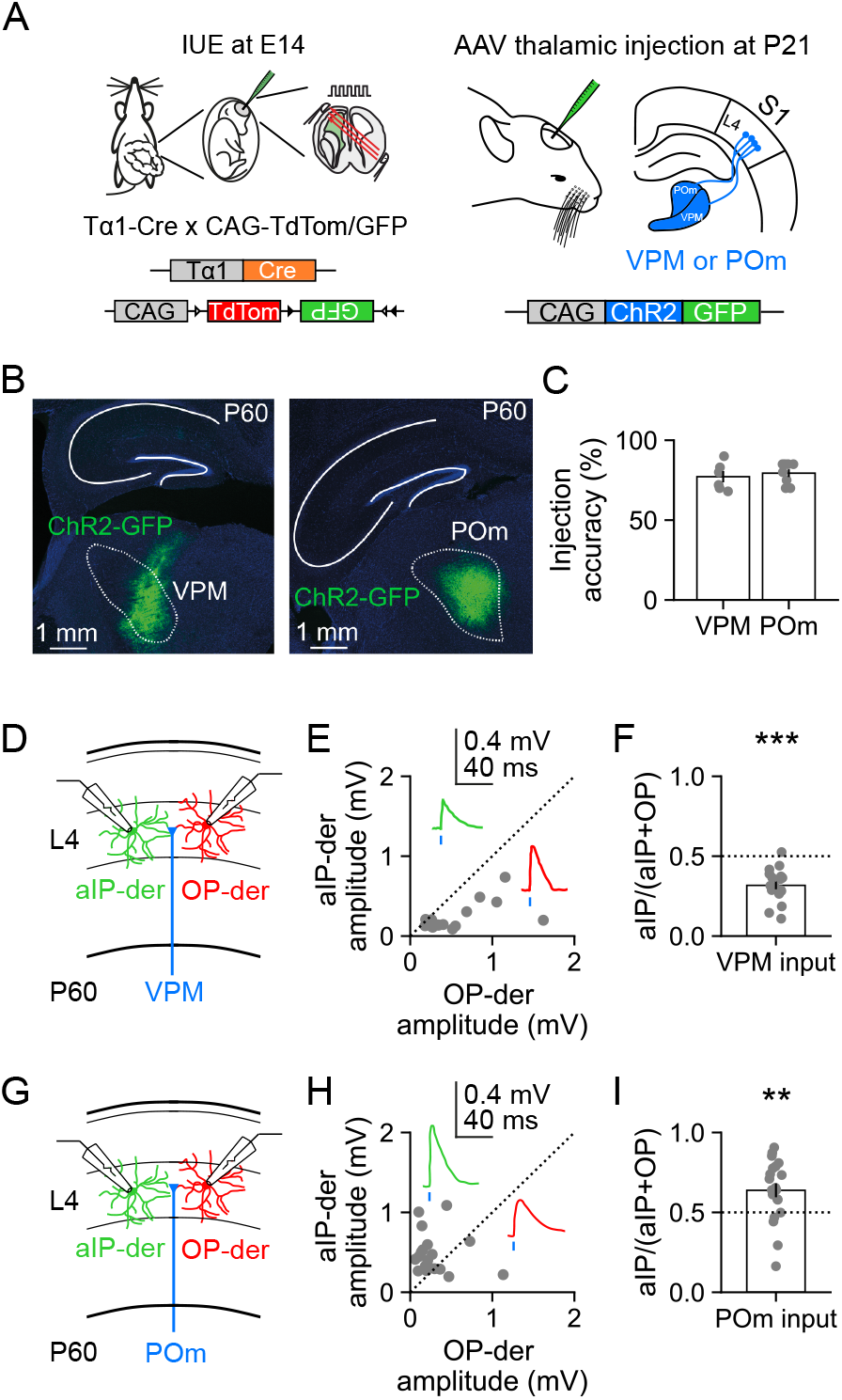
aIP-derived L4 neurons receive greater input from higher-order thalamus. (**A**) Experimental design for studying thalamic input to lineage-defined L4 neurons. IUE of a T*α*1-Cre and two-colour reporter plasmid was used to label aIP-derived (GFP^+^, green) and OP-derived (tdTomato^+^, red) L4 neurons in S1 (left). At P21, the mice received a thalamic injection of an AAV encoding CAG-ChR2-GFP, into either VPM or POm (right). (**B**) Histological analysis at P60 confirmed ChR2-GFP expression in the target thalamic nucleus. (**C**) Injection accuracy was defined as percentage ChR2-GFP expression restricted to the target nucleus. (**D**) To measure VPM input, simultaneous whole-cell recordings were performed from neuronal pairs comprising an aIP-derived and an OP-derived L4 neuron in acute slices, whilst ChR2-GFP^+^ VPM axons were stimulated with light pulses. (**E**) EPSP peak amplitudes for pairs of aIP-derived and OP-derived neurons in response to light stimulation of VPM axons. (**F**) aIP-derived neurons received weaker VPM input than OP-derived neurons (n = 18; p < 0.001, one sample t-test). (**G**) To measure POm input, simultaneous recordings were performed from an aIP-derived and an OP-derived L4 neuron, whilst ChR2-GFP^+^ POm axons were stimulated. (**H**) EPSP peak amplitudes for pairs of aIP-derived and OP-derived neurons in response to light stimulation of POm axons. (**I**) POm input was biased towards aIP-derived neurons, which received stronger POm input than OP-derived neurons (n = 21; p = 0.005, one sample t-test). Data represented as mean ± SEM, n = neuron pairs.

Activation of VPM axons (**Figure 4D**) elicited excitatory postsynaptic potentials (EPSPs) with short onset latencies, consistent with monosynaptic inputs to both populations of L4 neurons (response delay 3.78 ± 0.33 ms in aIP-derived neurons and 3.87 ± 0.40 ms in OP-derived neurons). However, the amplitude of the EPSP in the OP-derived neuron was consistently larger than in the paired aIP-derived neuron (**Figure 4E**), such that an index capturing the relative VPM input strength to aIP-derived neurons was significantly below 0.5 (**Figure 4F**; 0.32 ± 0.02). Meanwhile, activation of POm axons (**Figure 4G**) also elicited EPSPs with short onset latencies, consistent with monosynaptic inputs to both L4 populations (response delay 3.24 ± 0.44 ms in aIP-derived neurons and 3.45 ± 0.51 ms in OP-derived neurons). In contrast to the VPM input however, POm input revealed an amplitude bias in the reverse direction, such that EPSPs were consistently larger in the aIP-derived neuron than in the paired OP-derived neuron (**Figure 4H**), and an index of the relative POm input strength to aIP-derived neurons was significantly above 0.5 (**Figure 4I**; 0.64 ± 0.04). Neither a VPM nor a POm input bias was observed when pairs of unlabelled control L4 neurons were recorded (**Extended Data Figure 4**). These data establish that aIP-derived L4 neurons receive weaker input from VPM and stronger input from POm, when compared to neighbouring OP-derived neurons. This represents a novel lineage-dependent mechanism through which aIP-derived L4 neurons receive more input from higher-order thalamus.

### Lhx2 expression in developing L4 neurons is lineage-dependent

The transcription factor Lhx2 has been implicated in the development of S1, such that reducing Lhx2 expression levels across cortex results in an increase in L4 dendrites that do not target a specific barrel ^44,45^. To compare Lhx2 expression between our L4 excitatory neuronal populations, we used the same IUE strategy to label aIP-derived and OP-derived neurons with GFP and tdTomato (**Figure 5A**), then combined this with quantitative immunohistochemistry at P7, a time point at which L4 circuitry is being established ^46^. Lhx2 was found to be differentially expressed by L4 neurons at this age, such that aIP-derived neurons exhibited significantly lower levels of Lhx2 than OP-derived neurons (**Figure 5B-C** and **Extended Data Figure 5**; see **Methods**; 0.35 ± 0.03). This level of endogenous Lhx2 expression in aIP-derived L4 neurons is consistent with evidence that low Lhx2 levels favour dendrites that do not target a specific barrel, but also branch into septal regions outside barrels ^45^.

**Figure 5:**
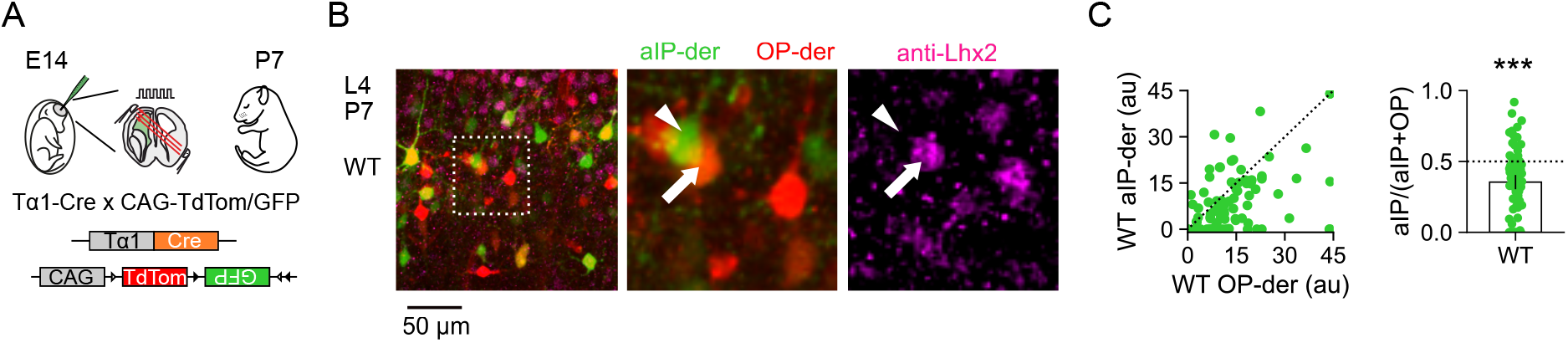
aIP-derived L4 neurons exhibit low levels of Lhx2 expression during postnatal development. (**A**) IUE of a T*α*1-Cre and two-colour reporter plasmid was used to label aIP-derived (GFP^+^, green) and OP-derived (tdTomato^+^, red) L4 neurons in S1. (**B**) Immunohistochemistry at P7 revealed lower levels of Lhx2 expression in aIP-derived neurons, compared to higher levels in OP-derived neurons. (**C**) Lhx2 expression levels were quantified (see **Methods**) in aIP-derived neurons and neighbouring OP-derived neurons within the same field of view (left; au, arbitrary units; values above 45 au have been set to 45 to aid visualisation). Normalized Lhx2 expression was significantly lower in aIP-derived neurons than OP-derived L4 neurons (right; n = 88, p < 0.001, one sample t-test). Data represented as mean ± SEM, n = neurons.

### Increased Lhx2 expression abolishes lineage-dependent differences in the sampling of thalamic inputs

To test whether Lhx2 expression is part of a lineage-based mechanism that determines how L4 neurons receive thalamic inputs, animals underwent IUE at E14 with the T*α*1-Cre, CAG-FLEx, plus an Lhx2 overexpression construct, CAG-Lhx2 (**Figure 6A** and **Extended Data Figure 6A**). This increased Lhx2 expression in aIP- derived neurons when compared to wild-type (WT) aIP-derived neurons at P7 (**Extended Data Figure 6B**), and normalised Lhx2 expression between the electroporated aIP-derived and OP-derived neurons within the same tissue (**Extended Data Figure 6C**). Compared to WT aIP-derived neurons, Lhx2-overexpressing (Lhx2) aIP- derived neurons occupied similar positions within S1 cortex (**Extended Data Figure 7A-C**), whilst *in vitro* recordings suggested that these neurons may exhibit subtle differences in their intrinsic electrical properties, but comparable levels of spontaneous synaptic activity (**Extended Data Figure 7D-J**). To characterise the dendritic morphology of Lhx2 aIP-derived L4 neurons, biocytin-filled neurons were reconstructed and co-registered to immunofluorescence images of the barrel field, as above (**Figure 6B-C**). The dendrites of Lhx2 aIP-derived L4 neurons overlapped significantly more with the principal barrel, and significantly less with areas outside barrels compared to WT aIP-derived L4 neurons (**Figure 6D**; WT overlap with principal barrel was 26.38 ± 4.03 %, adjacent barrel was 7.67 ± 1.91 %, and area outside barrels was 65.96 ± 4.31 %; Lhx2 overlap with principal barrel was 55.63 ± 7.81 %, adjacent barrel was 9.10 ± 3.25 %, and area outside barrels was 47.98 ± 7.88 %). This supports the idea that Lhx2 is part of a lineage-based mechanism by which L4 neurons acquire different thalamic inputs through their dendritic morphologies.

**Figure 6:**
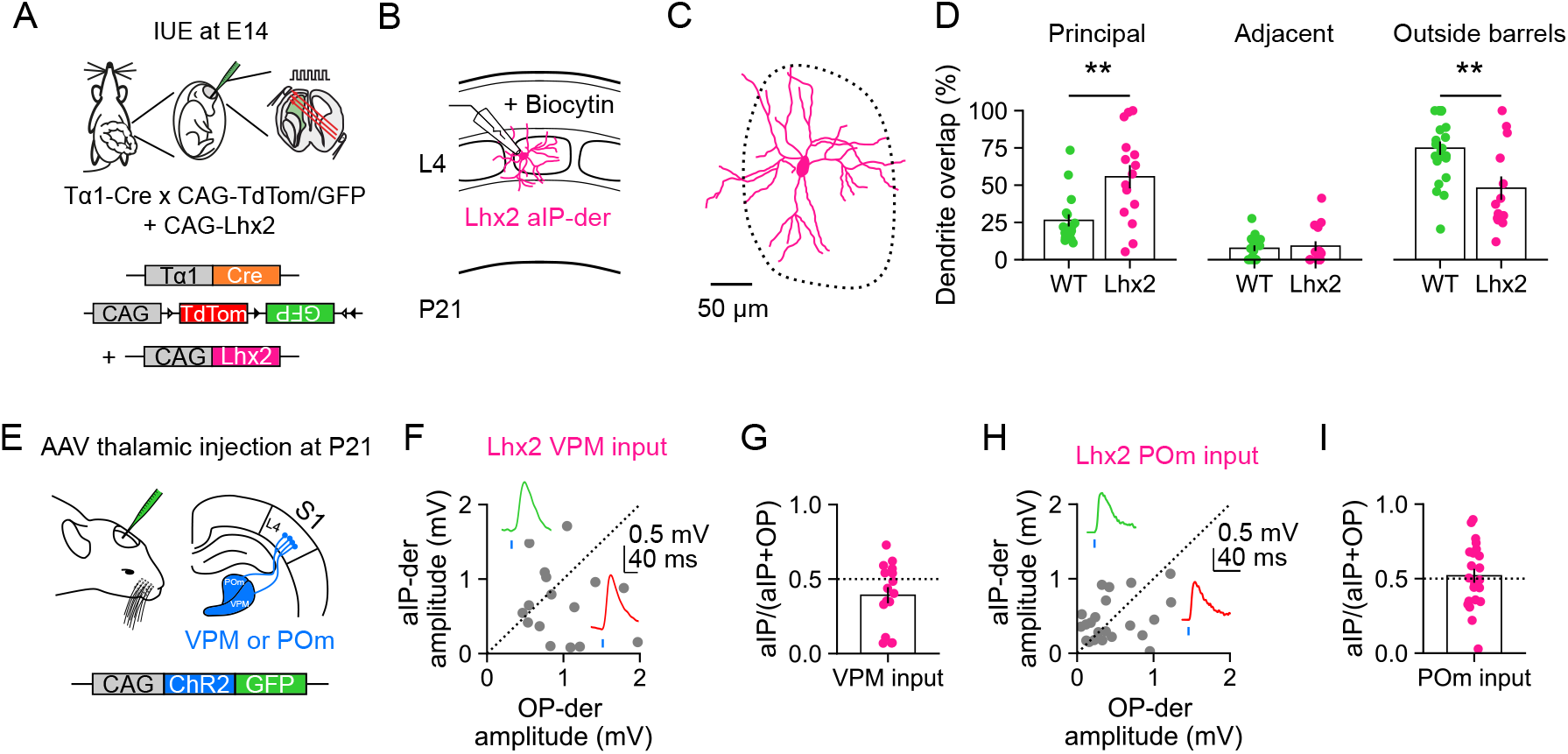
Increased Lhx2 levels in aIP-derived L4 neurons disrupts their higher-order thalamic input. (**A**) IUE of a T*α*1-Cre and reporter plasmid to label aIP-derived (GFP^+^, green) and OP-derived (tdTomato^+^, red) L4 neurons, was combined with a CAG-Lhx2 plasmid to increase Lhx2 expression levels. (**B**) Biocytin fills at P21 were used to reconstruct the dendritic morphology of aIP-derived L4 neurons overexpressing Lhx2 (Lhx2 aIP-derived). (**C**) Example reconstruction of an Lhx2 aIP-derived L4 neuron. (**D**) Dendrites of Lhx2 aIP-derived L4 neurons were more likely to target the principal barrel compared to WT (left; WT data from Figure 3; n = 17 and 15; p = 0.005, Mann-Whitney U test). Dendrites of both populations exhibited a similar degree of overlap with the adjacent barrel (middle; n = 17 and 15; p = 0.727, Mann-Whitney U test). Dendrites of Lhx2 aIP-derived L4 neurons were less likely to target the area outside of barrels, including septa, compared to WT (right; n = 17 and 15; p = 0.009, Mann-Whitney U test). (**E**) To study thalamic inputs, IUE was performed as in A, then mice received a thalamic injection at P21 of an AAV encoding CAG-ChR2-GFP, into either VPM or POm. (**F**) EPSP peak amplitudes for pairs of Lhx2-overexpressing aIP-derived and OP-derived neurons in response to light stimulation of VPM axons. (**G**) No statistically significant bias was detected in the strength of VPM input (n = 18; p = 0.06, one sample t-test). **(H**) EPSP peak amplitudes for pairs of Lhx2-overexpressing aIP-derived and OP-derived neurons in response to light stimulation of POm axons. (**I**) No statistically significant bias was detected in the strength of POm input (n = 22; p = 0.687, one sample t-test). Data represented as mean ± SEM, n = neurons/neuron pairs.

**Figure 7:**
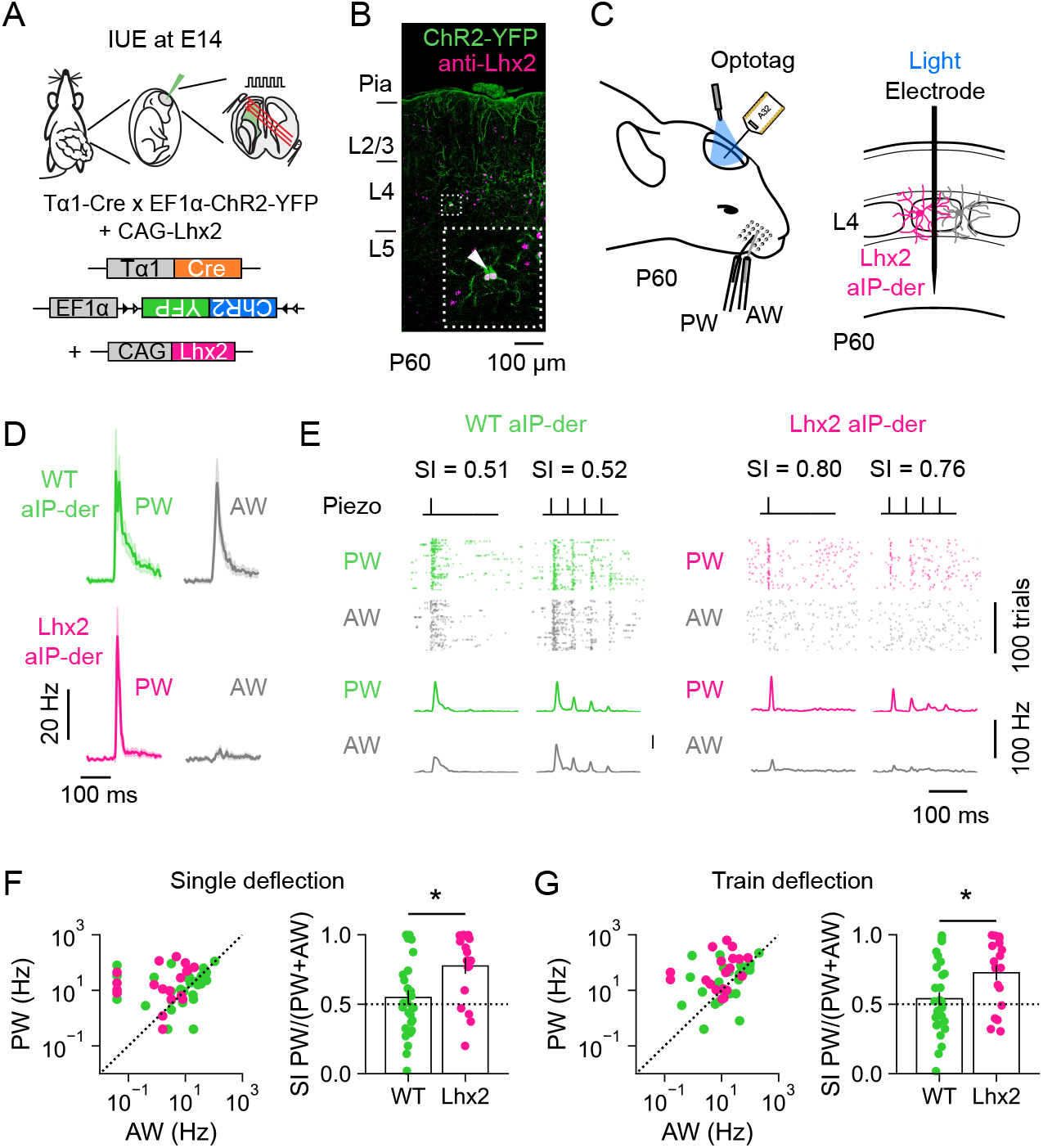
Increased Lhx2 levels disrupt multi-whisker response properties of aIP-derived L4 neurons. (**A**) IUE of a T*α*1-Cre, floxed ChR2-YFP and CAG-Lhx2 plasmid was used to optotag Lhx2-overexpressing aIP- derived L4 neurons in S1. (**B**) ChR2-YFP^+^ Lhx2 aIP-derived L4 neurons at P60. (**C**) The spiking activity of optotagged Lhx2 aIP-derived L4 neurons was recorded in response to the deflection of the PW or AW. (**D**) Mean responses of WT aIP-derived L4 neurons (data from Figure 2) and Lhx2 aIP-derived L4 neurons following a single deflection of the PW (left) or AW (right). (**E**) Spiking of an individual WT aIP-derived neuron (left) and an Lhx2 aIP-derived neuron (right) over 100 trials of either a single deflection (inner left) or trains of deflection (inner right) of the PW or AW. (**F**) Responses of individual WT and Lhx2 aIP-derived L4 neurons to single deflection of the PW and AW (left), and the distribution of corresponding SI values (right). Compared to WT, Lhx2 aIP-derived L4 neurons showed greater selectivity for the PW and relatively less responsivity to the AW (n = 29, 19; p = 0.010, Mann Whitney U test). (**G**) Responses to train deflection of the PW and AW (left), and corresponding SI values (right). Compared to WT, Lhx2 aIP-derived L4 neurons showed greater selectivity for the PW and relatively less responsivity to the AW (n = 29, 19; p = 0.024, Mann Whitney U test). Data represented as mean ± SEM, n = neurons, conventions as in Figure 2.

To test this prediction, we used the same IUE strategy at E14, before subsequently performing a thalamic injection of an AAV expressing ChR2-GFP into either VPM or POm at P21 (**Figure 6E**). Recordings from pairs of Lhx2 aIP-derived and OP-derived L4 neurons were then combined with light activation of ChR2-expressing thalamic axons in acute brain slices at P60. Activation of VPM or POm axons elicited EPSPs with short onset latencies (VPM response delay was 3.85 ± 0.35 ms and 3.4 ± 0.33 ms in Lhx2 aIP-derived and OP-derived; POm response delay was 3.75 ± 0.47 ms and 3.88 ± 0.46 ms in Lhx2 aIP-derived and OP-derived, respectively). However, unlike in the WT condition (**Figure 4**), EPSP amplitudes were comparable between aIP-derived L4 neurons and OP-derived neurons, suggesting that the sampling of thalamic inputs was no longer biased under conditions of increased Lhx2 overexpression. VPM-evoked EPSP amplitudes were not different for paired Lhx2 aIP-derived and OP-derived L4 neurons (**Figure 6F-G**; bias index of 0.53 ± 0.04) and POm-evoked EPSP amplitudes were not different for paired Lhx2 aIP-derived and OP-derived L4 neurons (**Figure 6H-I**; bias index of 0.52 ± 0.05). Taken together, these data suggest that increasing Lhx2 expression during development abolishes the lineage-dependent sampling of thalamic inputs by L4 neurons. For aIP-derived L4 neurons, this is characterised by a reduction in the relative levels of higher-order thalamic inputs from POm.

### Increased Lhx2 expression abolishes lineage-dependent higher-order response properties

To establish the functional consequences of the relative reduction in higher-order POm input to Lhx2 aIP-derived L4 neurons, we investigated the response properties of these neurons. Animals underwent IUE at E14 with T*α*1-Cre, floxed ChR2-YFP and CAG-Lhx2, to enable individual Lhx2 aIP-derived L4 neurons to be optotagged *in vivo* once the animals had reached P60 (**Figure 7A-C**). As above, extracellular recordings were performed whilst deflecting either the PW or AW (**Figure 7C**). PW deflection evoked robust responses in Lhx2 aIP-derived L4 neurons, whereas responses to AW deflection were greatly reduced (**Figure 7D-E**). As such, Lhx2 aIP-derived L4 neurons exhibited significantly higher SI values than WT aIP-derived L4 neurons in response to both single and train deflections of the whiskers (**Figure 7F-G**; single deflection mean SI was 0.55 ± 0.05 for WT and 0.76 ± 0.02 for Lhx2 aIP-derived neurons; train deflection mean SI was 0.55 ± 0.05 for WT and 0.78 ± 0.06 for Lhx2 aIP-derived neurons). Therefore, increased Lhx2 expression disrupts the multi-whisker response properties that are normally associated with aIP-derived L4 neurons, consistent with a reduction in how these neurons sample higher-order thalamic inputs.

### Increased Lhx2 expression disrupts sensory-evoked cortical plasticity

Higher-order thalamic inputs have been shown to promote sensory-evoked cortical plasticity through their ability to modulate the processing of first-order information ^47,48^. In rodent S1 for example, sensory-evoked plasticity between L4 and L2/3 is regulated by POm inputs ^14,16^. Given that Lhx2 influences the lineage-dependent sampling of higher-order thalamic inputs, we were keen to test whether this also affects sensory-evoked plasticity. We compared the WT condition in which animals underwent IUE with T*α*1-Cre and floxed ChR2-YFP at E14 to target L4 excitatory neurons, with the Lhx2 condition in which animals underwent IUE with T*α*1-Cre, floxed ChR2-YFP and CAG-Lhx2 (**Figure 8A**). Once the animals had reached P28, we performed extracellular recordings of multi-unit spiking activity in L2/3 of S1 (**Figure 8B**). ChR2-YFP expression enabled us to use robust light-evoked responses as confirmation that recordings targeted regions of S1 with high numbers of electroporated L4 neurons, which was further confirmed by histological analysis. A rhythmic whisker stimulation (RWS) protocol was used to induce sensory-evoked long term potentiation (sLTP; **Figure 8B**; see **Methods**)^14^. This involved measuring responses to multi-whisker deflections during a pre-induction period (Pre; 0.1 Hz deflections, 100 trials), then an sLTP induction period consisting of RWS at 8 Hz for 1 min, and finally a post-induction period (Post; 0.1 Hz deflection, 100 trials). In the WT condition, RWS induced robust sLTP of whisker-evoked responses in L2/3 (**Figure 8C-D**; pre WT spike rate was 43.29 ± 12.14 Hz and post spike rate was 137.06 ± 26.41 Hz), consistent with previous work ^14^. In the Lhx2 condition however, the same RWS protocol failed to elicit sLTP in L2/3 (**Figure 8E-G**; pre Lhx2 spike rate was 73.88 ± 30.10 Hz and post spike rate was 78.01 ± 31.73 Hz). This difference was not associated with differences in spiking levels during the RWS (**Extended Data Figure 8**), and control experiments confirmed that RWS was required to induce sLTP (**Figure 8H**; delta spike rate in WT without RWS was 106.49 ± 11.64 %, in WT with RWS was 338.59 ± 46.23 %, and in Lhx2 with RWS was 120.71 % ± 14.91). These data are consistent with a model in which higher-order thalamic inputs are required to elicit sensory-evoked plasticity in S1 ^14,16^, and demonstrate the functional importance of a lineage-dependent mechanism through which thalamic inputs are integrated in cortex.

**Figure 8:**
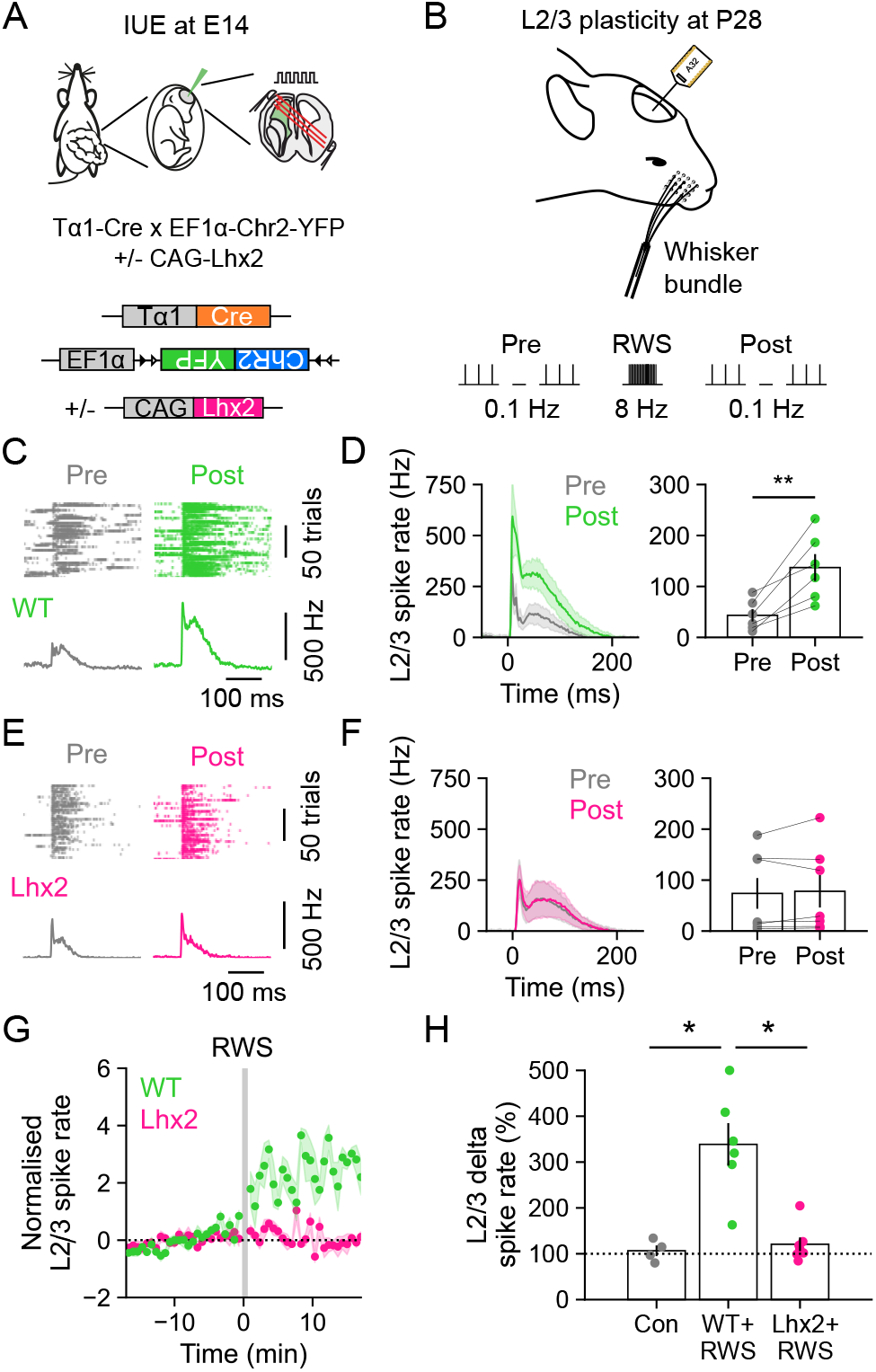
Higher-order input to aIP-derived L4 neurons is required for sensory-evoked plasticity in L2/3. (**A**) WT aIP-derived L4 neurons had undergone IUE of a T*α*1-Cre and floxed ChR2-YFP plasmid. Lhx2 aIP-derived L4 neurons had undergone IUE of the same plasmids, plus a CAG-Lhx2 plasmid. (**B**) Sensory-evoked plasticity was examined in L2/3 of S1 at P28 using a rhythmic whisker stimulation protocol (RWS; 8 Hz for 60 s). The presence of electroporated L4 neurons at the recording site was confirmed by ChR2 stimulation and subsequent histology. (**C**) Raster plots and PSTHs show multiunit L2/3 spiking activity in a WT animal. Responses to whisker deflections (0.1 Hz) are shown before (Pre) and after (Post) RWS. (**D**) Averaged (left) and separate (right) population data from WT animals reveal that RWS potentiated L2/3 activity (n = 6; p = 0.006, paired t-test). (**E**) Multiunit L2/3 activity in an Lhx2 animal. (**F**) Lhx2 animals did not exhibit potentiation of L2/3 activity following RWS (n = 7; p = 0.547, paired t-test). (**G**) Normalised L2/3 multiunit activity relative to the time of RWS (each data point is mean of five whisker deflections, 0.1 Hz). Shading indicates SEM around the mean. (**H**) Delta spike rate in WT animals that did not experience the RWS protocol (WT control), WT animals that experienced RWS (WT+RWS), or Lhx2 animals that experienced RWS (Lhx2+RWS) (n = 5, 6 and 7; p = 0.007, Kruskall Wallis; WT control vs. WT+RWS, p = 0.016; WT+RWS vs. Lhx2+RWS, p = 0.017, WT control vs. Lhx2+RWS, p = 1.00, Dunn’s test). Data represented as mean ± SEM, n = animals.

## Discussion

Here we combined *in utero* labelling, functional anatomy, and *in vivo* electrophysiology to investigate how distinct embryonic progenitor pools generate the cortical circuits that integrate thalamic inputs. Within mouse S1, L4 excitatory neurons derived from apical intermediate progenitors (aIP-derived) were compared to neighbouring neurons derived from other progenitors (OP-derived), at the same stage of embryonic development. Postnatal aIP-derived L4 neurons were found to exhibit features consistent with the integration of higher-order thalamic inputs, including multi-whisker responses, how their dendrites relate to the distribution thalamic axons, and their sampling of synaptic input from the higher-order thalamic nucleus, POm. Moreover, aIP-derived L4 neurons were shown to express low levels of the transcription factor Lhx2 during development relative to OP-derived neurons. Increasing Lhx2 in aIP-derived L4 neurons was sufficient to abolish their higher-order properties and disrupt the induction of sensory-evoked plasticity *in vivo*. These results identify a lineage-dependent mechanism through which cortical circuits are organised to receive thalamic inputs.

In agreement with earlier work in S1, L4 neurons exhibited heterogeneity in their spiking response properties following deflections of the PW and AW ^17,33,49,50^. By identifying L4 excitatory cortical neurons as a function of the embryonic progenitor type from which they derive, we were able to demonstrate that this heterogeneity is reflective of lineage. Previous evidence has shown that clonally-related excitatory neurons derived from the same individual progenitor cell can exhibit similar stimulus selectivity in the visual system ^51–53^. Our findings advance this observation by revealing that shared response properties are not only a feature of an individual clone, but can be a general feature of the neurons derived from particular types of excitatory embryonic progenitors. As a neuron’s spiking activity reflects its synaptic input, the differences in whisker-evoked responses were predicted to reflect differences in the inputs to the neuronal populations. We focused our investigation on the contribution of thalamic inputs, in line with evidence that response properties of S1 neurons can differ depending on the degree of input that they receive from higher-order thalamus ^13,15,54^. By combining optogenetic thalamic circuit mapping with paired recordings in L4, we found that aIP-derived neurons receive greater levels of input from POm and lower levels of input from VPM, consistent with the idea that the multi-whisker responses of the aIP-derived neurons reflect a greater targeting by higher-order thalamus ^4,32,54^. These data reinforce the notion that thalamic inputs from POm to S1 show heterogeneity at a cellular level ^24–26^ and reveal that lineage is a critical determinant of how individual cortical neurons sample inputs from thalamus.

Our morphological studies provided a potential explanation for the differential sampling of thalamic inputs to S1, whose anatomical organisation has been well characterised. Reconstructions of progenitor-defined L4 neurons revealed that whilst their soma positions were comparable, the dendrites of individual aIP-derived neurons were more likely to project outside of the principal barrel, when compared to OP-derived L4 neurons. This compliments previous evidence that L4 neurons vary in the degree to which their dendrites target barrels, and that neuronal morphology may be a mechanism through which progenitor-related properties are expressed ^40^. It will be interesting to explore whether morphological diversity represents a general mechanism through which progenitors influence the synaptic connectivity of their progeny. This could include future investigations into the relationship between the fine-scale morphology and functional connectivity of neurons derived from other progenitor types, and also how morphology relates to differences in intracortical connectivity, which may also contribute to lineage-dependent response properties ^29^.

To further explore the underlying mechanisms, we investigated the transcription factor Lhx2, which has been directly implicated in establishing the dendritic morphology of L4 excitatory neurons in S1 ^44,45^. We discovered that Lhx2 expression varies in a manner that reflects a cell’s lineage, and molecularly distinguishes L4 neurons, which have been characterised as a relatively homogenous neuronal population in the mature brain ^29,55^. During the period of postnatal cortical development, aIP-derived L4 neurons exhibited lower Lhx2 protein levels than OP-derived L4 neurons, consistent with a model in which low Lhx2 is associated with neurons that exhibit greater dendritic projections outside of the principal barrel ^45^. Experimentally increasing Lhx2 in aIP-derived L4 neurons resulted in dendrites that no longer projected outside of the principal barrel, abolished WT differences in how L4 neurons sampled inputs from higher-order POm and first-order VPM, and resulted in aIP-derived neurons with reduced multi-whisker response properties. These data implicate Lhx2 as part of a molecular mechanism within L4 cortical neurons, which establishes the lineage-dependent integration of thalamic inputs. This extends our understanding of how Lhx2 contributes to the coordination of fate specification and circuit assembly during the development of S1 ^44,45,56^, and supports the idea that Lhx2 can be used as a molecular tool to distinguish excitatory cortical subpopulations ^57^. Interestingly, the fact that Lhx2 has also been linked to activity-dependent genes ^45^, suggests a potential point of convergence through which progenitor-based mechanisms could interact with activity-dependent processes during circuit formation.

The manipulation of Lhx2 also allowed us to investigate the functional consequences of disrupting the lineage-dependent integration of thalamic inputs. Previous studies have shown that POm inputs to superficial layers of S1 are necessary for the induction of sLTP in L2/3 pyramidal neurons ^14^ and that combined L4 and POm activation is sufficient to induce LTP in brain slices ^16^. These findings have led to the proposal that the convergence of higher-order activity from POm, with first-order activity from VPM via L4, forms the basis of sLTP in L2/3 ^47,48^. Our experiments provide further support for this model by indicating that POm inputs to aIP-derived neurons within L4 are associated with a reduced ability to induce sLTP in L2/3 of S1. The data suggest that sensory-evoked plasticity requires that complementary information is delivered to different neuronal compartments and/or cortical layers. More broadly, our findings indicate that the cortical circuits underlying the integration of higher-order and first-order activity are the result of developmentally distinct lineages.

In summary, we establish that the integration of different thalamic inputs reflects the lineage of cortical neurons. The fact that thalamic inputs exhibit lineage dependency raises the possibility that common developmental principles govern long-range and local connectivity within cortex ^28–30,58^, in order to generate functional subnetworks. These observations suggest that the evolution of multiple progenitor types does not only serve to expand cortical volume or increase the representation of particular post-mitotic cell types, but also to generate distinct routes of information flow through cortex.

## Data availability

All data and reagents detailed in this paper are available from the corresponding author upon reasonable request.

## Author contributions

M.J.B. and C.J.A. designed the study. M.J.B., K.M., J.J.R., G.G., S.V.A. and T.J.E. conducted electrophysiology experiments. M.J.B., K.M. and G.G. performed anatomical experiments. M.J.B., K.M., J.J.R. and G.G. wrote software and performed analysis. S.E.N. generated molecular tools. M.J.B. and C.J.A. wrote the paper.

## Acknowledgements

We would like to thank members of the Akerman lab, Adam Packer, Zoltan Molnar and Louise Upton for advice and comments. Tarik Haydar and Ulrich Müller generously provided reagents, and Adam Packer generously provided access to Neurolucida software. The research leading to these results has received funding from the European Research Council under grant agreement number 617670; plus BBSRC project BB/S007938/1. In addition, M.J.B. was supported by a University of Oxford Clarendon Scholarship, G.G. and S.V.A. by a Wellcome Trust Doctoral Fellowship, T.J.E by a MRC Career Development Award, and S.E.N. by a Royal Society Dorothy Hodgkin Fellowship.

## Methods

### Mice

Experiments were performed using both male and female C57/BL6 wildtype mice, which were bred, housed, and used in accordance with the UK animals (Scientific Procedures) Act (1986). Breeding females were checked for plugs daily and the day of plugging was considered embryonic day (E) 0.5.

### In utero electroporation

*In utero* electroporation (IUE) was performed using standard procedures at E14 consistent with previous studies targeting S1 L4 ^34,35^. Briefly, pregnant females were anaesthetised using isofluorane (Zoetis). Buprenorphine (Vetergesic; 0.1 mg/kg) and meloxicam (Metacam; 5mg/kg) were administered subcutaneously. The uterine horns were exposed by midline laparotomy. A mixture of plasmid DNA (∼2 µg/µl) and 0.03 % fast green dye (Sigma Aldrich) was injected intraventricularly using micropipettes pulled from borosilicate glass capillaries (1.5 mm outer diameter, Warner Instruments), through the uterine wall and amniotic sac. Plasmid DNA included: Tα1-Cre, in which Cre recombinase is under the control of a portion of the Tubulin alpha-1 (Tα1) promoter ^37^; CAG-tdTomato/GFP, which uses the chicken β-actin (CAG) promoter to control a flexible excision (FLEx) cassette, whereby Cre recombination permanently switches expression from tdTomato to enhanced green fluorescent protein (GFP) ^43^; EF1α-ChR2-YFP (pAAV-EF1a-double floxed-hChR2(H134R)-EYFP-WPRE-HGHpA; Addgene #20298; http://n2t.net/addgene:20298; RRID:Addgene_20298; a gift from Karl Deisseroth), in which Cre recombination turns on the expression of channelrhodopsin-2 fused to enhanced yellow fluorescent protein (ChR2-EYFP) under the control of the human elongation factor-1a promoter; CAG-Lhx2 (pAAV-CAG- Lhx2), in which mouse Lhx2 is under the control of the CAG promoter. To generate CAG-Lhx2, mouse Lhx2 cDNA was amplified from TetO-FUW-Lhx2 (Addgene #61537; http://n2t.net/addgene:61537; RRID:Addgene_61537; a gift from Rudolf Jaenisch) and cloned into the AAV backbone derived from pAAVCAG- iCre (Addgene #51904; http://n2t.net/addgene:51904 ; RRID:Addgene_51904; a gift from Jinhyun Kim). Plasmids were injected as a 1:1 ratio and the total volume injected per embryo was ∼2 µl. The anode of a 5 mm Platinum Tweezertrode (BTX) was placed over the dorsal telencephalon outside the uterine muscle. Five pulses (50 ms duration separated by 950 ms) at 36 V were delivered with an ECM 830 pulse generator (BTX). The uterine horns were placed back inside the abdomen, the cavity filled with warm physiological saline, and the abdominal muscle and skin incisions were closed with Vicryl (Ethicon) and Prolene (Ethicon) sutures, respectively. Dams were monitored until the birth of the pups and further analgesia was provided, as appropriate.

### Postnatal intrathalamic viral injections

Animals that had undergone IUE were used for targeted intracerebral injections at P21. Briefly, mice were anaesthetised using isofluorane and placed in a stereotaxic frame (Kopf Instruments). Vetergesic (0.1 mg/kg) was administered subcutaneously, and EMLA cream (Aspen) was applied to the scalp. An incision was made to expose the skull. Bregma and lamda were located and a small craniotomy was performed to expose the neocortex. Injections were targeted to either the ventral posteromedial nucleus (VPM) (1.8 mm lateral to bregma, 1.4 mm posterior; 3.1 mm deep from pia), or the posterior medial nucleus (POm) (1.4 mm lateral to bregma, 2.1 mm posterior; 3 mm deep from pia) of the thalamus. 120-240 nl of an adeno-associated virus (AAV) carrying CAG-ChR2-GFP, in which ChR2-GFP was under the control of the CAG promoter (Boyden, UNC Vector Core), was injected over a period of 8 minutes using a pulled glass micropipette (Blaubrand intraMARK). The craniotomy was covered, and the skin closed with Vicryl sutures. Further analgesia was provided, as appropriate. *Post hoc* histological analysis was used to confirm correct location of the thalamic injections.

### In vitro slice preparation and recording conditions

Acute slices were prepared from postnatal animals from P21 (range P21 – 28), or from P60 (range P60 – 75) where a postnatal intracerebral injection had been performed. Animals were anaesthetised using isofluorane and decapitated. Thalamocortical 350-400 µm slices (55 ° with respect to midline) were cut using a vibrating microtome (Microm). Slices were prepared in artificial cerebrospinal fluid (aCSF) containing (in mM): 65 sucrose, 85 NaCl, 2.5 KCl, 1.25 NaH_2_PO_4_, 7 MgCl_2_, 0.5 CaCl_2_, 25 NaHCO_3_ and 10 glucose, pH 7.2-7.4, bubbled with carbogen gas (95% O_2_ / 5% CO_2_). Slices were immediately transferred to a storage chamber containing aCSF (in mM): 130 NaCl, 3.5 KCl, 1.2 NaH_2_PO_4_, 2 MgCl_2_, 2 CaCl_2_, 24 NaHCO_3_ and 10 glucose, pH 7.2 - 7.4, at 32°C, and bubbled with carbogen gas. When required, slices were transferred to a recording chamber and continuously superfused with aCSF bubbled with carbogen gas with the same composition as the storage solution (32 °C and perfusion speed of 2 ml/min). Whole-cell current-clamp recordings were performed using glass pipettes, pulled from borosilicate glass capillaries (1.2 mm outer diameter, Warner Instruments), containing (in mM): 110 potassium gluconate, 40 HEPES, 2 ATP-Mg, 0.3 Na-GTP, 4 NaCl and 4 mg/ml biocytin (pH 7.2-7.3; osmolarity 290-300 mosmol/l).

### In vitro stimulation, recording protocols and analysis

Recordings were made using a Multiclamp 700B (Molecular Devices) amplifier and acquired using WinWCP (University of Strathclyde, UK) or pClamp (Molecular Devices) software. All recordings were low pass filtered at 2 kHz and digitized at a sampling frequency of 10 kHz. Slices were placed into a recording chamber and barrels were visualised in layer 4 (L4) under brightfield illumination. The distribution of cells labelled by IUE meant that the intrinsic electrophysiological properties and morphology of neurons were sampled across the extent of S1. Single L4 excitatory neurons within barrels were identified and targeted using video assisted Dodt contrast imaging. Progenitor identity was confirmed using fluorescent light. The intrinsic properties of the recorded neurons were assessed using a variety of protocols consisting of hyperpolarising and depolarising current steps (from -300 to +600 pA, 100 pA steps) in current clamp. Measurements included resting membrane potential, spike threshold, spike frequency, spike amplitude and inter-spike interval . The resting membrane potential was calculated from a pre-stimulus period of 0 pA current injection, averaged over 10 sweeps. The values of spike threshold voltage were calculated manually from the recorded traces. All other measures were calculated using custom scripts written in python. Spontaneous excitatory postsynaptic currents (sEPSCs) were recorded in voltage clamp whilst holding the cell at -70 mV. Each sEPSC recording was 10 minutes long. The pClamp ‘event detection’ tool was used to create a standardised template by manually selecting ∼200 spontaneous events. This template was then used to automatically detect sEPCSs. The amplitude and frequency of the sEPCSs were calculated using custom scripts written in MATLAB. Monosynaptic thalamic inputs to layer 4 (L4) neurons were studied by stimulating ChR2-GFP expressing axons in L4, which originated from either POm or VPM. Photoactivation of ChR2 was achieved using 1 ms light pulses via an LED (473 nm; 3.8-21.6 mW/mm^2^, LedEngin) and the amplitude of short-latency, time-locked, light-evoked excitatory postsynaptic potentials (EPSPs) were measured from pairs of simultaneously recorded L4 neurons from the average of 10-40 sweeps. Light intensity was adjusted to produce low amplitude monosynaptic EPSPs (mean peak <3 mV), to minimize the chance of recruiting polysynaptic activity.

### In vivo recording conditions

Extracellular recordings were performed from P60 (range P60 – 75) for selectivity experiments, and from P28 (range P28 – 35) for plasticity experiments. Animals were anaesthetised with 25 % urethane (1 g/kg; Sigma) in phosphate-buffered saline (PBS; Thermo Fisher), then mounted in a stereotaxic frame (Stoelting) and continuously supplied with oxygen (0.3 ml/min) throughout the recording. Glycoperronium Bromide (Glycopyrrolate; 0.01 mg/kg) was administered subcutaneously, and Marcaine (Aspen) was applied to the scalp. A heat mat controlled by a direct current temperature regulation system (FHC inc.) was used to maintain body temperature at 37 °C. A single incision was made to remove the skin from the skull and a craniotomy of ∼2 mm diameter was performed using a pneumatic dental drill (Foredom). A 32-channel single-shank electrode (Neuronexus) was repeatedly submerged in 1,1’-Dioctadecyl-3,3,3’,3’-Tetramethylindocarbocyanine Perchlorate (DiI) lipophilic dye (2.5 mg/ml, in 70 % ethanol, Thermo Fisher) and then slowly inserted into the cortex at an angle of 20 ° from the vertical (with respect to bregma: 3 mm lateral,1.2 mm posterior; 0.9 mm deep from pia).

### In vivo stimulation and recording protocols

The electrode was connected to an acquisition board (OpenEphys) using a NPD36 connector (Omnetics) and a RHD2132 amplifier (Intan). In order to deflect two single whiskers independently, borosilicate capillaries (1.5 mm outer diameter, Warner Instruments) were attached to two piezoelectric bending actuators (Piezo Technics). Deflection was achieved using a single 100 Hz sinusoidal waveform controlled by a Piezo Controller (Piezo Technics). Photoactivation of ChR2 was achieved using 10 ms light pulses via an LED (473 nm, 45 mW/mm^2^, LedEngin) positioned above the cortical surface. All stimulus protocols were generated using custom scripts written in MATLAB and delivered via a PulsePal pulse train generator (OpenEphys). The principal whisker (PW) relative to the electrode insertion site was manually identified online by the presence of a robust, short-latency spiking response following whisker deflection and a characteristic current source density profile (CSD), calculated as the second spatial derivative of the local field potential (LFP). The adjacent whisker (AW) was defined as the whisker immediately rostral to the principal whisker. If this whisker was missing or did not evoke a response, the whisker immediately caudal was used. The identities of the PW and AW were confirmed offline using the population spike rate and response latency of all L4 excitatory neurons from a given animal.

To study sensory long-term potentiation (sLTP), we performed a rhythmic whisker stimulation (RWS) protocol as has been described previously (Gambino et al., 2014). Before (‘pre’) and after (‘post’) RWS, responses were measured by deflecting multiple whiskers (∼10) with a single piezoelectric bending actuator at 0.1 Hz for 100 trials. During RWS, the whiskers were deflected for 1 min at 8 Hz (i.e., 100 Hz waveform every 125 ms). In all experiments, baseline activity was recorded for between 30 min and 1 hr following electrode insertion. Electrodes were immersed in 1% Tergazyme (Sigma Aldrich) for 2 hr between recording sessions. Data was acquired at 30 kHz. To obtain multiunit activity (MUA), data was bandpass filtered between 300-6000 Hz. MUA was detected using the median absolute deviation of the filtered signal ^59^. To obtain the LFP, data was lowpass filtered under 300 Hz and a 50 Hz notch filter was applied. To obtain single unit activity, data was spike sorted using Kilosort ^31^, and curated in Phy ^60^.

### In vivo analysis

Cortical layers were defined using the PW CSD. The shortest latency CSD sink evoked upon stimulation of the principal whisker was defined as L4. Electrode channels above L4 were assigned as L2/3. Regular spiking (primarily excitatory) neurons were separated from fast-spiking (putative interneurons) neurons using a spike waveform trough-to-peak time of 0.5 ms as the separation criterion. To identify neurons that were directly activated by ChR2, repeated light pulses were delivered to the probe insertion site. Optotagged neurons were defined by a mean response latency < 5 ms. To control for the recruitment of polysynaptic light-evoked activity, the AMPA receptor blocker DNQX (10 mM, Tocris) was applied to the cortical surface in a subset of experiments. The stimulus-evoked response window for single whisker deflection was defined as 50 ms. In all experiments, spontaneous activity was calculated on a trial-by-trial basis as the average spike rate of a given unit or channel and subtracted from stimulus-evoked responses. Response latency was calculated as the 1 ms time bin containing the first spike 5 – 20 ms following whisker deflection in each trial. The selectivity index for individual neurons in response to single deflection was calculated as: RPW / RPW + RAW, where RPW and RAW were the cumulative spontaneous subtracted spike count in the 50 ms following deflection of the PW or AW, respectively. For train deflection RPW and RAW were the total sum of the spike counts in four separate 50 ms windows following each deflection. The stimulus-evoked response window for the deflection of multiple whiskers (i.e., **Figure 8**) was defined as 250 ms.

### Histological analysis

Following whole-cell patch-clamp recording, acute brain slices were fixed in 4 % paraformaldehyde (PFA, Sigma Aldrich) in 0.1 M PBS (pH 7.4). Biocytin-filled cells were visualised using repeated incubation with streptavidin alexa fluor 680 (1:1000, Thermo Fisher), boosted using anti-streptavidin (1:1000, goat, Vector Laboratories). For whole-brain histology, brains were fixed by cardiac perfusion of PBS followed by 4 % PFA. Brains were stored in 4 % PFA for an additional 24 hr, after which they were washed and stored in PBS. Embryonic brains were fixed by submersion in 4 % PFA for three days, after which they were washed in PBS and mounted in 4 % agar (Intralabs) to aid sectioning. Tissue was sectioned at 50 µm on a vibrating microtome (Microm).

The standard immunohistochemistry protocol was as follows. Sections were washed three times in PBS for 5 min, then blocked in 20 % normal goat serum (NGS, Sigma Aldrich) in 0.1% Triton-X (Thermo Fisher) in PBS (PBST) for 2 h at RT. Sections were washed in PBS and incubated overnight with primary antibody diluted in 0.1 % PBST at 4 °C. Primary antibodies included: anti-VGlut2 (1:250, rabbit, Synaptic Systems), anti-GFP (1:1000, chicken, Aves Lab), anti-RFP (1:1000, rat, Chromotek), and anti-Lhx2 (1:500, rabbit, Abcam). VGLUT2 staining was facilitated using heated antigen retrieval at 92 °C in 10 mM fresh sodium citrate (pH 6.0) for 30 min prior to primary antibody incubation. Slices were washed in PBS and were incubated for 2 hr with secondary antibodies diluted in 0.1 % PBST at RT. Secondary antibodies included: anti-chicken alexa fluor 488 (1:1000, goat, Thermo Fisher), anti-rat alexa fluor 568 (1:1000, goat; Thermo Fisher), and anti-rabbit alexa fluor 635 (1:1000, goat, Thermo Fisher). Sections were counter-stained with 4’,6-Diamidino-2-Phenylindole, Dihydrochloride (DAPI, 1:10000, Thermo Fisher) and mounted on slides with VectaShield (Vectorlabs). Fluorescent images were acquired with a LSM 880 confocal microscope (Zeiss) equipped with 488 nm, 561 nm, and 633 nm lasers and a 20x water-immersion objective (W Plan-Apochromat) using ZEN software (Zeiss). All cell counting, localisation and fluorescence analysis was performed using FIJI (ImageJ).

Soma position was analysed with custom scripts written in MATLAB. Soma coordinates were labelled manually, and a reference line drawn along the L4/L5 boundary based on VGLUT2 fluorescence. The image and soma coordinates were then transformed with respect to the reference line, thereby straightening the cortical layers whilst maintaining relative soma position. Barrel boundaries were detected automatically based on VGLUT2 fluorescence in L4. All soma were assigned a barrel index score with 1 indicating a soma located in the middle of a barrel, and 0 indicating a soma located at the midpoint between two barrel boundaries. Biocytin-filled neurons were reconstructed using Neurolucida and Neuroexplorer software (MBF Bioscience) and co-registered to immunofluorescence images of the barrel field. The identity of the principal barrel was defined anatomically as the barrel with which the majority of dendrites overlapped. The adjacent barrel was defined as the next closest barrel. Dendritic overlap was quantified as a percentage of total dendritic length. Neurons with < 5 % total dendritic length within any barrel were excluded from further analyses. Lhx2 expression levels were quantified as a ratio of the mean pixel intensity (MPI) of the cell body of an aIP-derived L4 neuron over the mean MPI of three neighbouring OP-derived L4 neurons within the same z-plane and within a 100 x 100 µm region of interest.

### Statistical analysis

All data are presented as mean ± standard error unless otherwise stated. All statistical analysis was performed in Python using SciPy. Continuous data were assessed for normality and appropriate parametric or non-parametric statistical tests were applied (* p < 0.05, ** p < 0.01, *** p < 0.001).

**Extended Data Figure 1:**
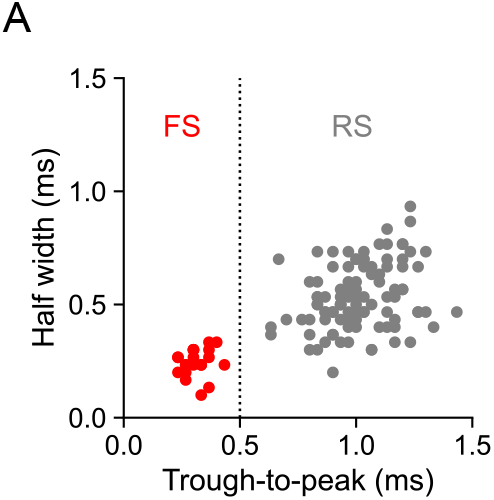
*In vivo* identification of putative excitatory neurons from extracted waveform properties. (**A**) Regular-spiking (RS; primarily excitatory) neurons were distinguished from fast-spiking (FS; primarily interneurons) neurons based on the waveform properties of single units isolated using Kilosort. A trough-to-peak time of 0.5 ms was used as the separation criterion (FS trough-to-peak was 0.31 ± 0.01 ms and RS trough-to-peak was 1.01 ± 0.02 ms, n = 21 and 100, respectively), in keeping with previous studies in rodent cortex ^61^. Data represented as mean ± SEM, n = neurons.

**Extended Data Figure 2:**
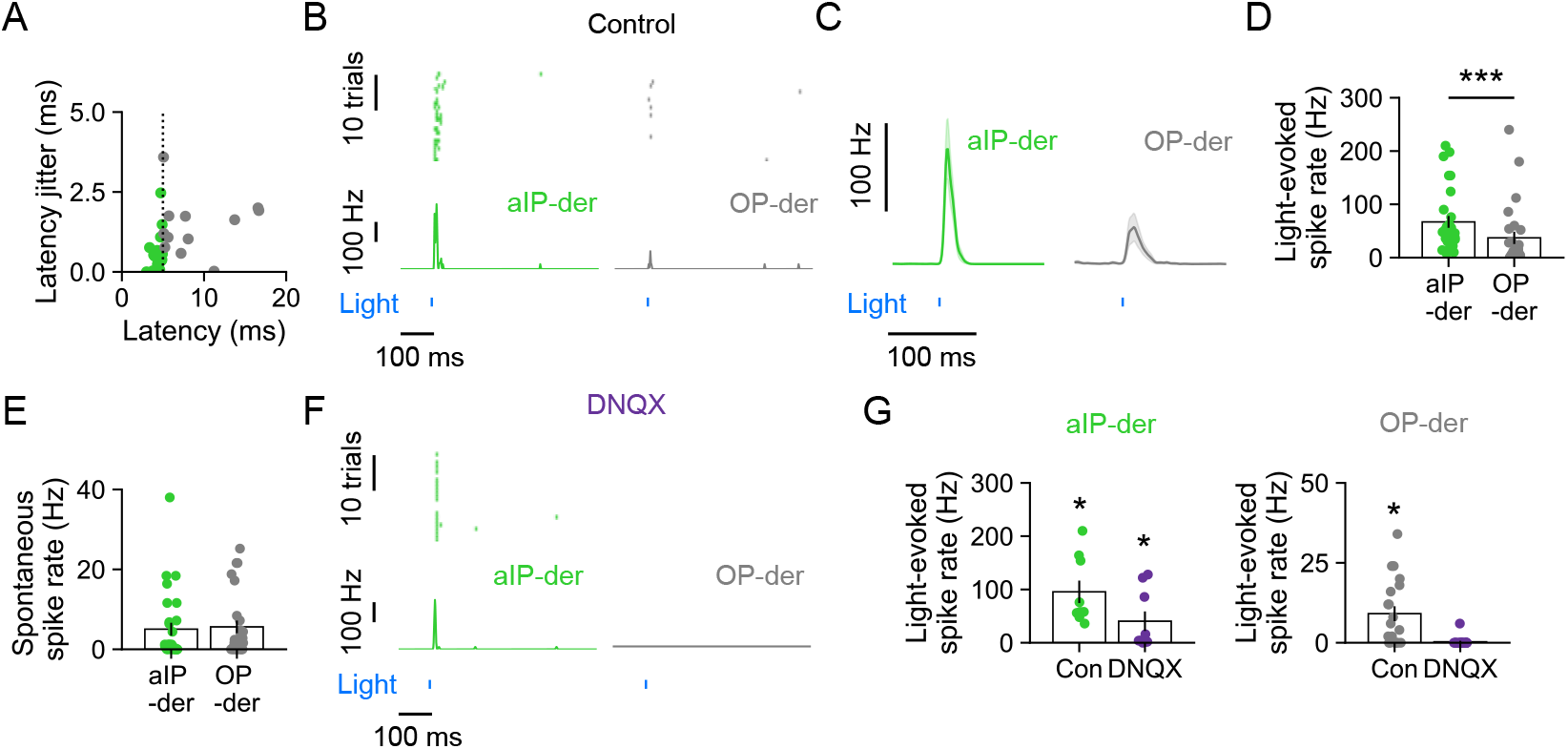
*In vivo* identification of aIP-derived L4 neurons by optotagging. (**A**) Optotagging was used to distinguish putative aIP-derived (i.e., ChR2-YFP^+^) L4 neurons from putative OP-derived L4 neurons. Based on spike times in response to light pulses (25 repeats, 10 ms pulse), a mean light-evoked spike latency of 5 ms was used as the separation criterion (aIP-derived spike latency was 4.34 ± 0.10 ms, OP-derived spike latency was 8.97 ± 1.18 ms, n = 29 and 26). (**B**) Light-evoked spiking of an individual aIP-derived neuron (left) and OP-derived neuron (right). (**C**) Mean light-evoked spiking response of a population of aIP-derived and OP-derived neurons (n = 29 and 26). (**D**) Light-evoked spiking was significantly higher in aIP-derived L4 neurons (aIP-derived spike rate was 66.97 ± 11.02 Hz, OP-derived was 37.08 ± 11.56 Hz; n = 29 and 26; p < 0.001, Mann Whitney U test). (**E**) Spontaneous spike rates did not differ between aIP-derived and OP-derived neurons (aIP-derived spontaneous spike rate was 5.03 ± 1.60 Hz, OP-derived was 5.62 ± 1.57 Hz; n = 29 and 26; p = 0.24, Mann Whitney U test). (**F**) Example recordings show that after blocking synaptic transmission with the glutamate receptor blocker DNQX, light-evoked spiking was still evident in an optotagged aIP-derived neuron but was abolished in a nearby OP-derived neuron. (**G**) Population data revealed that DNQX reduced, but did not abolish, light-evoked responses in aIP-derived neurons (control was 95.56 ± 21.01 Hz, DNQX was 40.44 ± 18.35 Hz; p = 0.004 and p = 0.012, one sample Wilcoxon against a median of zero; n = 9). On the other hand, DNQX did abolish light-evoked responses in OP-derived neurons (control was 9.10 ± 2.32 Hz, DNQX was 0.30 ± 0.30 Hz; p = 0.001, p = 0.317, one sample Wilcoxon against a median of zero; n = 20). Data represented as mean ± SEM, n = neurons.

**Extended Data Figure 3:**
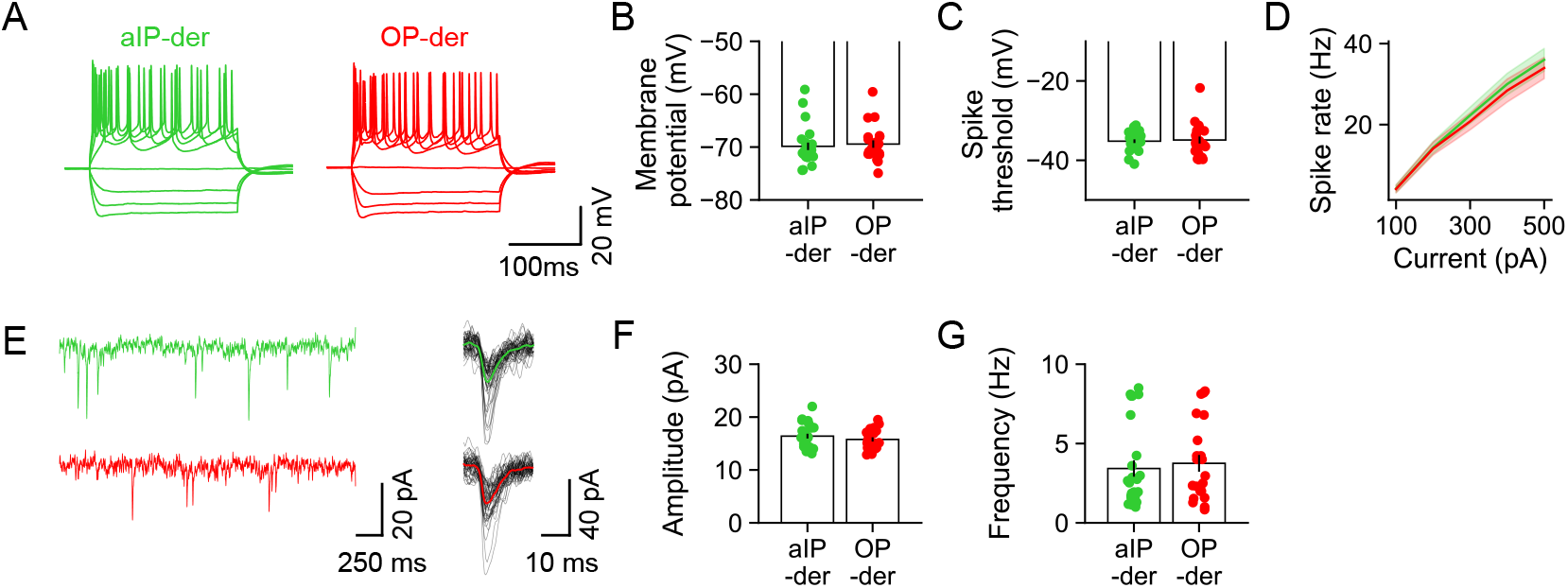
Intrinsic electrical and synaptic properties of aIP-derived L4 neurons. (**A**) Current clamp recordings performed in acute brain slices from an aIP-derived and OP-derived L4 neuron, in response to current steps. (**B**) Resting membrane potential did not differ between the aIP-derived and OP-derived L4 neurons (aIP-derived was -67.74 ± 1.16 mV, OP-derived was -70.76 ± 0.99 mV; n = 19 and 14; p = 0.07, t-test). (**C**) Spike threshold did not differ between the aIP-derived and OP-derived L4 neurons (aIP-derived was -35.18 ± 0.46 mV, OP-derived was -34.89 ± 0.88 mV; n = 25 and 22; p = 0.44, Mann Whitney U test). (**D**) Spike rate in response to injected current did not differ between the aIP-derived and OP-derived L4 neurons (100, 200, 300, 400 and 500 pA steps; aIP-derived was 4.20 ± 0.98 Hz, 14.30 ± 1.49 Hz, 22.30 ± 2.01 Hz, 30.00 ± 2.65 Hz, and 36.00 ± 2.80 Hz; OP-derived was 4.09 ± 0.90 Hz, 14.09 ± 1.58 Hz, 20.79 ± 2.08 Hz, 28.41 ± 2.85 Hz, 33.98 ± 2.62 Hz; n = 25 and 22; p = 0.26, Mixed ANOVA). (**E**) Voltage clamp recordings of spontaneous excitatory synaptic currents in an aIP-derived and an OP-derived L4 neuron. Composite traces represent the mean of 50 spontaneous events. (**F**) The amplitude of spontaneous excitatory synaptic currents did not differ between aIP-derived and OP-derived L4 neurons (aIP-derived was 16.40 ± 0.45 pA, OP-derived was 15.77 ± 0.40 pA; n = 26 and 22; p = 0.31, t-test). (**G**) The frequency of spontaneous excitatory synaptic currents did not differ between aIP-derived and OP-derived L4 neurons (aIP-derived was 3.55 ± 0.49 Hz, OP-derived was 3.79 ± 0.39 Hz; n=21 and 19; p = 0.28, Mann Whitney U test). Data represented as mean ± SEM, n = neurons.

**Extended Data Figure 4:**
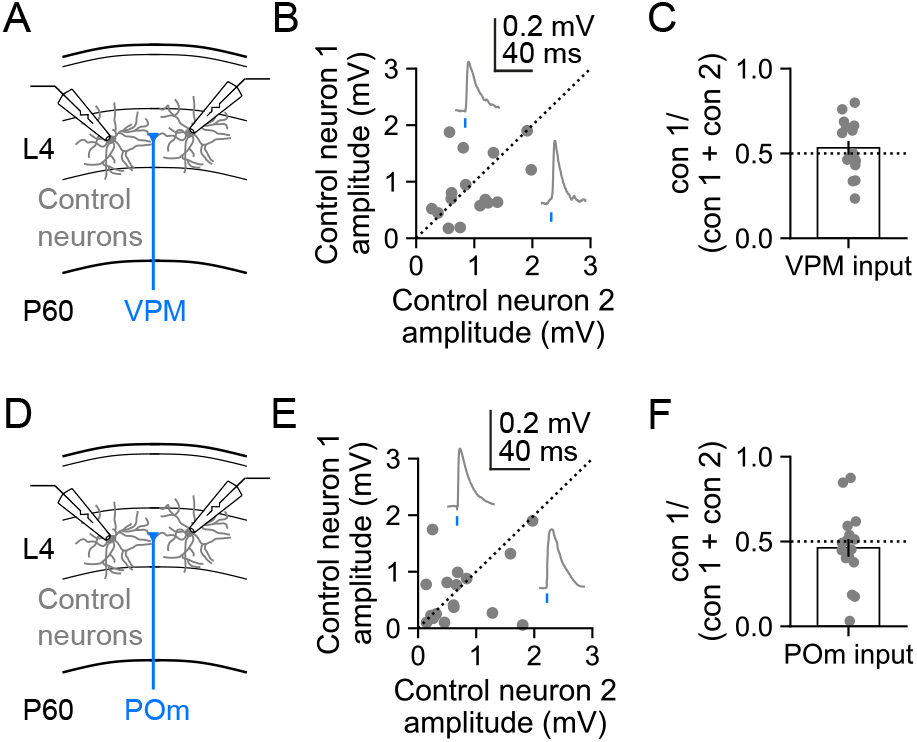
No consistent bias of thalamic inputs is observed in pairs of unlabelled L4 neurons. (**A**) Simultaneous whole cell recordings were performed from pairs of unlabelled control L4 neurons at P60, whilst ChR2-GFP^+^ VPM axons were stimulated with light pulses. (**B**) EPSP peak amplitudes for pairs of unlabelled control neurons in response to light stimulation of VPM axons. (**C**) No consistent bias in VPM input was observed under these conditions (0.53 ± 0.04; n = 18; p = 0.428, one sample t-test). (**D**) In a separate set of experiments, simultaneous whole cell recordings were performed from pairs of unlabelled control L4 neurons, whilst ChR2-GFP^+^ POm axons were stimulated. (**E**) EPSP peak amplitudes for pairs of unlabelled control neurons in response to light stimulation of POm axons. (**F**) No consistent bias in POm input was observed under these conditions (0.46 ± 0.05; n = 18; p = 0.4, one sample t-test). Data represented as mean ± SEM, n = neuron pairs.

**Extended Data Figure 5:**
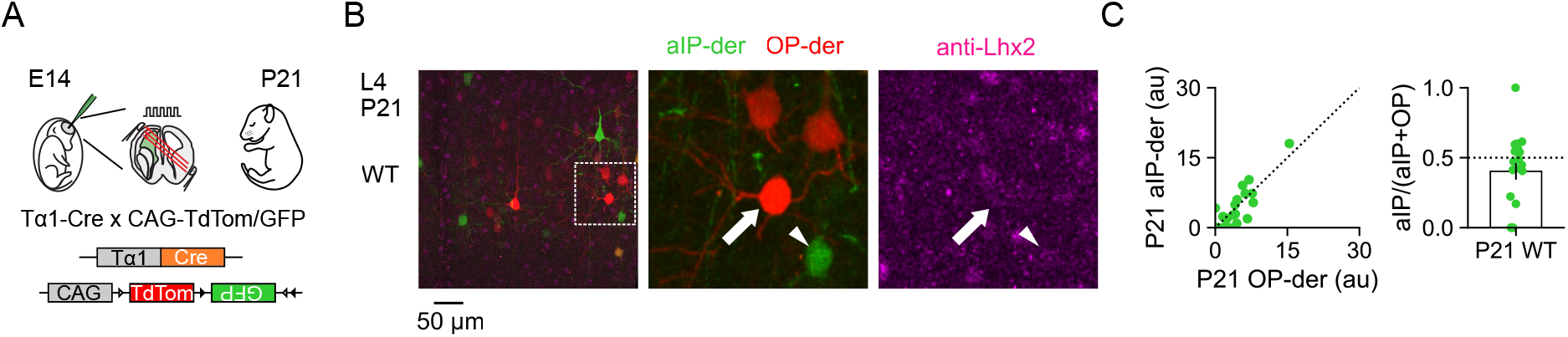
Lhx2 expression has decreased and does not show lineage-dependent differences by P21. (**A**) At E14, animals underwent IUE of a T*α*1-Cre and two-colour reporter plasmid to label aIP-derived (GFP^+^, green) and OP-derived (tdTomato^+^, red) L4 neurons in S1. (**B**) Immunohistochemistry at P21 revealed low levels of Lhx2 expression in both aIP-derived and OP-derived neurons (contrast levels enhanced to aid visualisation). (**C**) Lhx2 expression levels were quantified in aIP-derived neurons and neighbouring OP-derived neurons within the same field of view (left; au, arbitrary units). Normalized Lhx2 expression was not different in aIP-derived and OP-derived L4 neurons at P21 (right; 0.40 ± 0.06; n = 20 and 20; p = 0.23, one sample Wilcoxon test against a median of 0.5). Data represented as mean ± SEM, n = neurons.

**Extended Data Figure 6:**
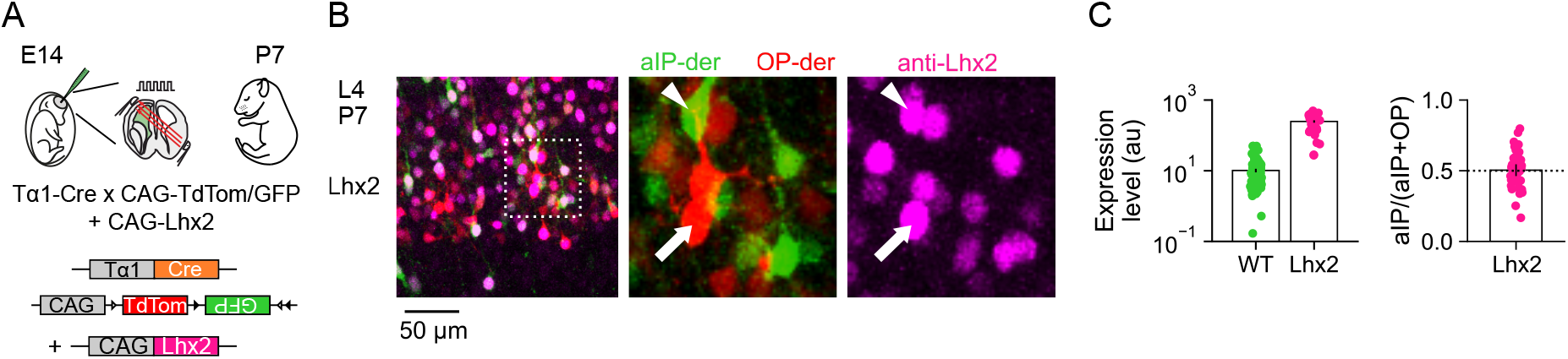
An overexpression construct increases Lhx2 levels in electroporated L4 neurons. (**A**) To raise Lhx2 levels in aIP-derived neurons, a CAG-Lhx2 overexpression plasmid was delivered with T*α*1-Cre and a two-colour reporter plasmid by IUE. (**B**) Immunohistochemistry at P7 revealed similarly high levels of Lhx2 expression in aIP-derived neurons and OP-derived electroporated neurons. (**C**) Quantification confirmed increased levels in Lhx2-overexpressing (Lhx2) aIP-derived L4 neurons compared to WT aIP-derived L4 neurons (left; WT expression was 10.18 au ± 1.16, Lhx2 expression was 248.57 ± 19.51). Within the Lhx2 electroporated tissue, expression levels were similar in the aIP-derived and OP-derived electroporated L4 neurons, such that there was no difference in relative expression (right; 0.50 ± 0.02; n = 40 and 40, p = 0.85, one sample t-test). Data represented as mean ± SEM, n = neurons.

**Extended Data Figure 7:**
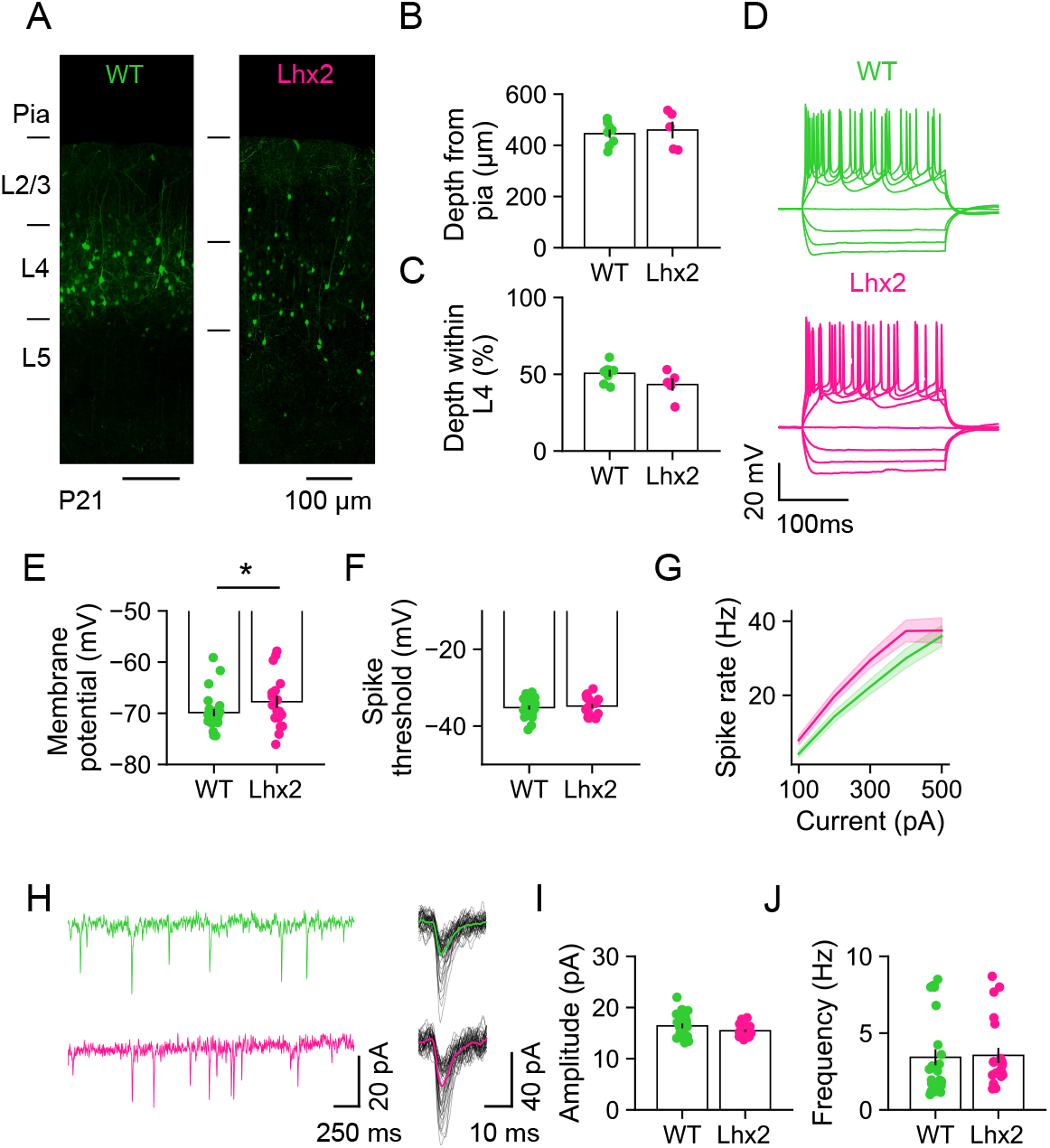
Effects of increased Lhx2 on migration, intrinsic electrical and synaptic properties of aIP-derived L4 neurons. (**A**) Wild-type aIP-derived neurons (WT; left) at P21, labelled via IUE of the T*α*1-Cre plasmid and a reporter plasmid at E14. Lhx2-overexpressing aIP-derived neurons (Lhx2; right) at P21, labelled via IUE of the T*α*1-Cre plasmid, a reporter plasmid and a CAG-Lhx2 plasmid to increase Lhx2 expression levels in the electroporated neurons. (**B**) The somata of WT and Lhx2 neurons were located at similar distances from the pia mater (WT depth of 445.67 ± 16.04 µm, Lhx2 depth of 459.77 ± 32.80 µm; n = 8 and 5; p = 0.67, t-test). (**C**) WT and Lhx2 neurons occupied similar depths within L4 (WT at 50.64 ± 2.14 %, Lhx2 at 43.32 ± 4.05 %; n = 8 and 5; p = 0.11, t-test). (**D**) Example current clamp recordings from a WT and Lhx2 aIP-derived L4 neuron in response to current steps. (**E**) Resting membrane potential was slightly more depolarised in Lhx2 aIP-derived neurons compared to WT (WT was -69.86 ± 0.71 mV, Lhx2 was -67.74 ± 1.16 mV; n = 25 and 19; p = 0.042, Mann Whitney U test). (**F**) Spike threshold was not statistically different between WT and Lhx2 aIP-derived neurons (WT was -35.18 ± 0.46 mV, Lhx2 was -34.60 ± 0.84 mV; n = 25 and 14; p = 0.51, t-test). (**G**) Spike rate in response to injected current was not statistically different between WT and Lhx2 aIP-derived neurons (100, 200, 300, 400 and 500 pA steps; WT was 4.20 ± 0.98 Hz, 14.30 ± 1.49 Hz, 22.30 ± 2.01 Hz, 30.00 ± 2.65 Hz, 36.00 ± 2.80 Hz; Lhx2 was 7.89 ± 1.27 Hz, 19.73 ± 1.67 Hz, 29.47 ± 2.09 Hz, 37.37 ± 2.95 Hz, 37.50 ± 3.39 Hz; n = 25 and 19; p = 0.067, Mixed ANOVA). (**H**) Example voltage clamp recordings of spontaneous excitatory synaptic currents in a WT and an Lhx2 aIP-derived L4 neuron. (**I**) The amplitude of spontaneous excitatory synaptic currents was not statistically different (WT was 16.40 ± 0.45 pA, Lhx2 was 15.47 ± 0.25 pA; n = 26 and 21; p = 0.09, t-test). (**J**) The frequency of spontaneous excitatory synaptic currents was not statistically different between WT and Lhx2 aIP-derived neurons (WT was 3.43 ± 0.52 Hz, Lhx2 was 3.55 ± 0.49 Hz; n = 26 and 21; p = 0.18, Mann Whitney U). Data represented as mean ± SEM, n = sections or neurons.

**Extended Data Figure 8:**
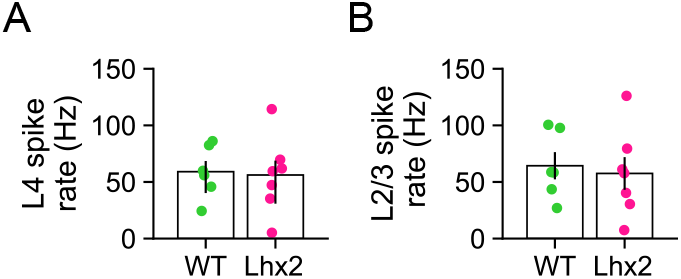
Overall spiking activity during the RWS protocol was comparable across conditions. (**A**) Multiunit spiking activity in L4 during the RWS (8 Hz for 60 s) was similar for animals with WT or Lhx2 aIP-derived L4 neurons (WT was 59.10 ± 9.43 Hz, Lhx2 was 56.17 ± 12.66 Hz; n = 6 and 7; p = 0.86, t-test). (**B**) Multiunit spiking activity in L2/3 during RWS was not different for animals with WT or Lhx2 aIP-derived L4 neurons (WT was 64.31 ± 12.00 Hz, Lhx2 was 57.53 ± 14.42; n = 6 and 7; p = 0.73, t-test). Data represented as mean ± SEM, n = animals.

